# CRISPRa-mediated disentanglement of the Dux-MERVL axis in the 2C-like state, totipotency and cell death

**DOI:** 10.1101/2024.11.25.625195

**Authors:** Paul Chammas, Sheila Q Xie, Lessly P Sepulveda-Rincon, Bryony J Leeke, Marian H Dore, Dirk Dormann, Ryan T Wagner, Michael T McManus, Mohammad M Karimi, George Young, Michelle Percharde

**Affiliations:** MRC Laboratory of Medical Sciences, London W12 0HS; Institute of Clinical Sciences, Imperial College, London W12 0HS; University of California at San Francisco, San Francisco, California 91413, USA; Department of Molecular Pharmacology and Experimental Therapeutics, Mayo Clinic, Rochester, MN 55901, USA; Comprehensive Cancer Centre, School of Cancer & Pharmaceutical Sciences, Faculty of Life Sciences & Medicine, King’s College London, London, United Kingdom

## Abstract

Transposable elements (TEs) provide sequences that are powerful cis-regulatory drivers of gene expression programmes. This is particularly apparent during early development when many TEs become de-repressed. MERVL elements are highly yet transiently upregulated in mouse totipotent 2-cell (2C) embryos during major zygotic genome activation (ZGA), and in 2C-like cells in vitro. One of the most powerful activators of MERVL is the pioneer transcription factor, Dux. However, apparent differences lie in the requirement for Dux versus MERVL activation in embryos, for unclear reasons. Moreover, sustained Dux activation causes cell toxicity in multiple cell types, which may or may not be linked to MERVL activation. Using a CRISPR-activation, 2C-GFP reporter system, we have unpicked the relative role of Dux and MERVL in ZGA, totipotent-like characteristics and cell toxicity. We find that direct MERVL activation comprises only a portion of the Dux-dependent transcriptome, and which is sufficient for expanded fate potential, but not other totipotency features. Conversely, Dux-induced pathology is independent of MERVL activation and involves induction of the pro-apoptotic factor, Noxa. Our study highlights the complexity of the Dux-MERVL transcriptional network and uncovers a new player in Dux-driven pathology.

## Introduction

Transposable elements (TEs) are repetitive, genomic sequences that are either historically or currently mobile within our DNA. Of these, endogenous retrovirus (ERV) sequences make up approximately 8-10% of mammalian genomes and have arisen from ancient viral infections. While TE activation can be harmful – driving DNA damage, mutagenesis or inflammation (Gasior et al. 2006; Malki et al. 2014; Burns 2017; De Cecco et al. 2019) – TE-mediated spread of cis-regulatory elements has played a powerful role in gene expression networks (Kunarso et al. 2010; Sundaram and Wysocka 2020). In particular, the Long Terminal Repeat (LTR) promoters of ERVs may harbour transcription factor binding sites and act as alternative promoters or enhancers for nearby genes (Wang et al. 2007; Chuong et al. 2013; Fuentes et al. 2018; Modzelewski et al. 2021).

Although repressed in most somatic contexts, specific families of TEs are upregulated in early embryos (Fadloun et al. 2013; Ohno et al. 2013; Jachowicz et al. 2017; Percharde et al. 2018). In mice, murine endogenous retrovirus-L (MERVL) elements are highly and specifically activated in 2-cell stage embryos, then rapidly repressed upon 2-cell exit (Kigami et al. 2003; Peaston et al. 2004; Svoboda et al. 2004; Macfarlan et al. 2011; Macfarlan et al. 2012). The MERVL LTR (MT2_Mm) acts as a powerful promoter for nearby genes and drives activation of 2-cell (2C) specific genes in embryos (Macfarlan et al. 2011; Kruse et al. 2019). The activation of MERVL or the 2C programme has been suggested to be essential for zygotic genome activation (ZGA) and embryo progression (Huang et al. 2017; Sakashita et al. 2023; Yang et al. 2024). Interestingly, transient MERVL activation is also seen in rare cells arising from cultures of embryonic stem cells (ESCs), which express 2C-specific genes and TEs (Macfarlan et al. 2012; Ishiuchi et al. 2015). Moreover, these cells, termed 2C-like cells, display other markers of 2-cell embryos such as increased chromatin accessibility and downregulation of pluripotency factor expression (Boskovic et al. 2014; Ishiuchi et al. 2015; Eckersley-Maslin et al. 2016; Kruse et al. 2019). They also show totipotent-like characteristics: an ability to contribute to extraembryonic lineages in chimeras (Macfarlan et al. 2012; Choi et al. 2017; Yang et al. 2020). These data point to an integral role for MERVL in the 2-cell embryo and the 2C-like state.

Despite this, the mechanism of MERVL upregulation, and its exact role in ZGA, is still unclear. In ESCs, the homeobox transcription factor Dux has been found to potently activate MERVL elements and induce the 2C-like state (De Iaco et al. 2017; Hendrickson et al. 2017; Whiddon et al. 2017). Interestingly, its human homolog, DUX4 also activates HERVL, MLT2A1 and MLT2A2 TEs during human ZGA at the 4-8 cell stage (Geng et al. 2012; Hendrickson et al. 2017). However *in vivo*, Dux knockout (KO) impedes development but is ultimately viable (Chen and Zhang 2019; Guo et al. 2019; De Iaco et al. 2020), with other factors presumably sustaining MERVL and 2C/ZGA gene expression (Gassler et al. 2022; Ji et al. 2023; Guo et al. 2024). Conversely, failure to shut down expression of Dux/DUX4 and TE expression in various cell and embryo models has been associated with DNA damage, cell death, or developmental arrest (Geng et al. 2012; Percharde et al. 2018; Guo et al. 2019; Olbrich et al. 2021; Xie et al. 2022). DUX4 de-repression in muscle cells moreover drives cell death and the human disease, Facioscapulohumeral muscular dystrophy (FSHD)(Dixit et al. 2007; Lemmers et al. 2010). Thus, Dux/DUX4 must be tightly controlled in normal cells. This raises the question – which parts of the Dux-MERVL axis are most important for development and 2-cell characteristics? And are the negative consequences of Dux overexpression due to MERVL activation or independent from TE upregulation?

Here, we assess both the beneficial and pathological contributions of Dux and MERVL activation in ESCs and embryos. With CRISPR-activation (CRISPRa) and our recently described 2C-GFP/CD4 reporter ESCs (Xie et al. 2022), we show that direct Dux activation much more faithfully resembles endogenous 2C-like cells than 2C-like cells driven by MERVL alone. However, despite MERVL only activating a portion of 2C-like genes and features, it is equally efficient in conferring expanded fate potential. Conversely Dux, but not MERVL, is associated with reduced proliferation and cell death, and involves activation of the pro-apoptotic factor, Noxa. Our study places MERVL as a partial yet key feature of the 2C-like state and reveals its separation from the pathological impacts of Dux over-expression.

## Results

### Generation of a 2C-GFP/CD4 CRISPRa system

To investigate the relationship between Dux and MERVL activation, we used a CRISPRa system combined with our recently published 2C-GFP/CD4 reporter mouse ESCs (Xie et al. 2022). Approximately 1-2% of ESCs enter transiently into a 2C-like state, termed 2C-like cells, which can be identified by MERVL-driven GFP expression and purified by FACS or CD4-based magnetic isolation. These ESCs were targeted with a dead Cas9 construct fused to the tripartite VP16-p16-RelA activator (dCas9-VPR (CV))(Chavez et al. 2015) into the *Rosa26* locus (Fig.1A, S1A-B). We confirmed the functionality of dCas9-VPR by transducing heterozygous *Rosa26*^*CV/+*^ ESCs with sgRNAs targeting two silenced genes, *Hbb-bh1* or *Ttn*, which led to high and specific activation of their relative targets (Fig.1B). To enable activation of endogenous MERVL loci, we continued with a single *Rosa26*^*CV/+*^, 2C-GFP/CD4 clone for all subsequent experiments (Fig. S1B), referred to subsequently as CVG ESCs. Three independent sgRNAs were selected that target the MERVL consensus sequence and were each predicted to match 65-70% of individual MERVL copies (Fig.1C, S1C and Table S1). Transient transfection of CVG ESCs with these guides alongside two Dux sgRNAs (Fig. S1D) or a negative control guide (sgGAL4) resulted in a robust increase in 2C-GFP-positive cells compared to the control (Fig.1D). Note that sgGAL4 is not predicted to bind to any mouse genes, so 2C-like cells arising from this transfection represent spontaneously occurring, or endogenous, 2C-like cells. We next validated upregulation of *Dux* or MERVL RNA by RT-qPCR. MERVL expression is induced in GFP-positive cells from all three sgRNA transfections (Fig.1E-F). In contrast, *Dux* is activated only in sgGAL4 and sgDux cells and not sgMERVL.

**Figure 1:**
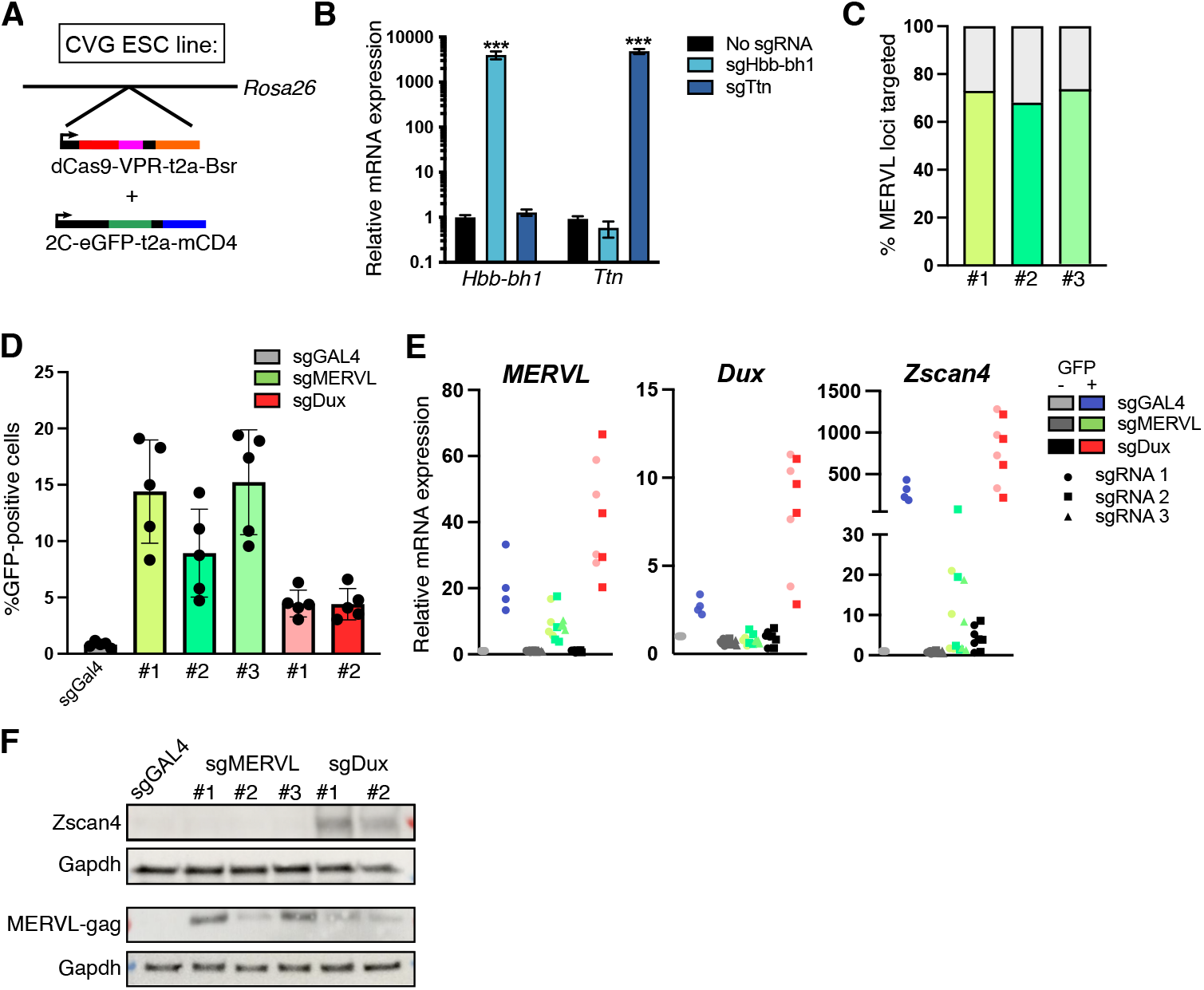
Establishment of a CRISPR-activation system for 2C-like cell generation. A) Diagram of CVG ESCs: a single clone of E14 ESCs targeted with dCas9-VPR into the *Rosa* locus then a 2C-GFP/CD4 construct by random integration. B)qRT-PCR validation of CRISPRa by dCas9-VPR in ESCs transduced with the indicated sgRNAs. Data are mean +/-s.e.m of 3 independent Rosa^dCas9-VPR/+^ clones. *** P <0.001, two-tailed student’s t-test. C)Predicted binding of the indicated sgRNAs to MERVL MT2_Mm loci. D)Flow cytometry analysis of the percentage of GFP-positive cells in CVG ESCs 48 h after transfection with the indicated sgRNAs. Data are mean +/-s.e.m of n=5 independent experiments. E)qRT-PCR analysis of the indicated genes in FACS-sorted GFP-negative (-) and positive (+) cells. Data represent n=4 independent experiments. F)Western blot in unsorted CVG ESCs 48 h after transfection with sgRNAs, representative of 3 independent experiments.

This is consistent with MERVL elements as downstream of Dux. *Zscan4* is interestingly not activated by sgMERVL, suggesting it to be a direct Dux target (Fig.1E). These data confirm the establishment of an efficient system to activate endogenous *Dux* or MERVL in ESCs, and begin to point to differences between the 2C-like cells they induce.

### MERVL CRISPRa generates an intermediate 2C-like transcriptome

We next set out to probe the global transcriptional impact of Dux and MERVL (Fig.2A). CVG ESCs were transfected with either Dux, MERVL, or GAL4 sgRNAs and pure populations of GFP-positive or - negative cells sorted 48 hours later and processed for RNA-seq (Fig.2A, Table S2). We first looked at the expression of TEs, with clustering of samples based on TE subfamily expression revealing a clear separation between 2C-GFP positive (2C-like) and 2C-GFP negative (ESC) samples (Fig.2B). Interestingly, while GAL4/Dux-induced 2C-like cells cluster together, furthest away from ESCs, MERVL-induced 2C-like cells exhibit an intermediate profile. We examined the top 10 most upregulated TE subfamilies in endogenous 2C-like cells (sgGAL4, GFP+). As expected, MERVL-int and its promoter, MT2_Mm are activated in all conditions, as well as other ERVL subfamilies such as MT2C_Mm and MLT2E (Fig.2C, S2A-B). In contrast, elements such as major satellites (GSAT_MM), and the ERV3 member MT2B1, some of which are also upregulated in 2-cell embryos (Fig. S2B), are not induced by sgMERVL. These data reveal only a partial TE activation profile upon MERVL CRISPRa.

**Figure 2:**
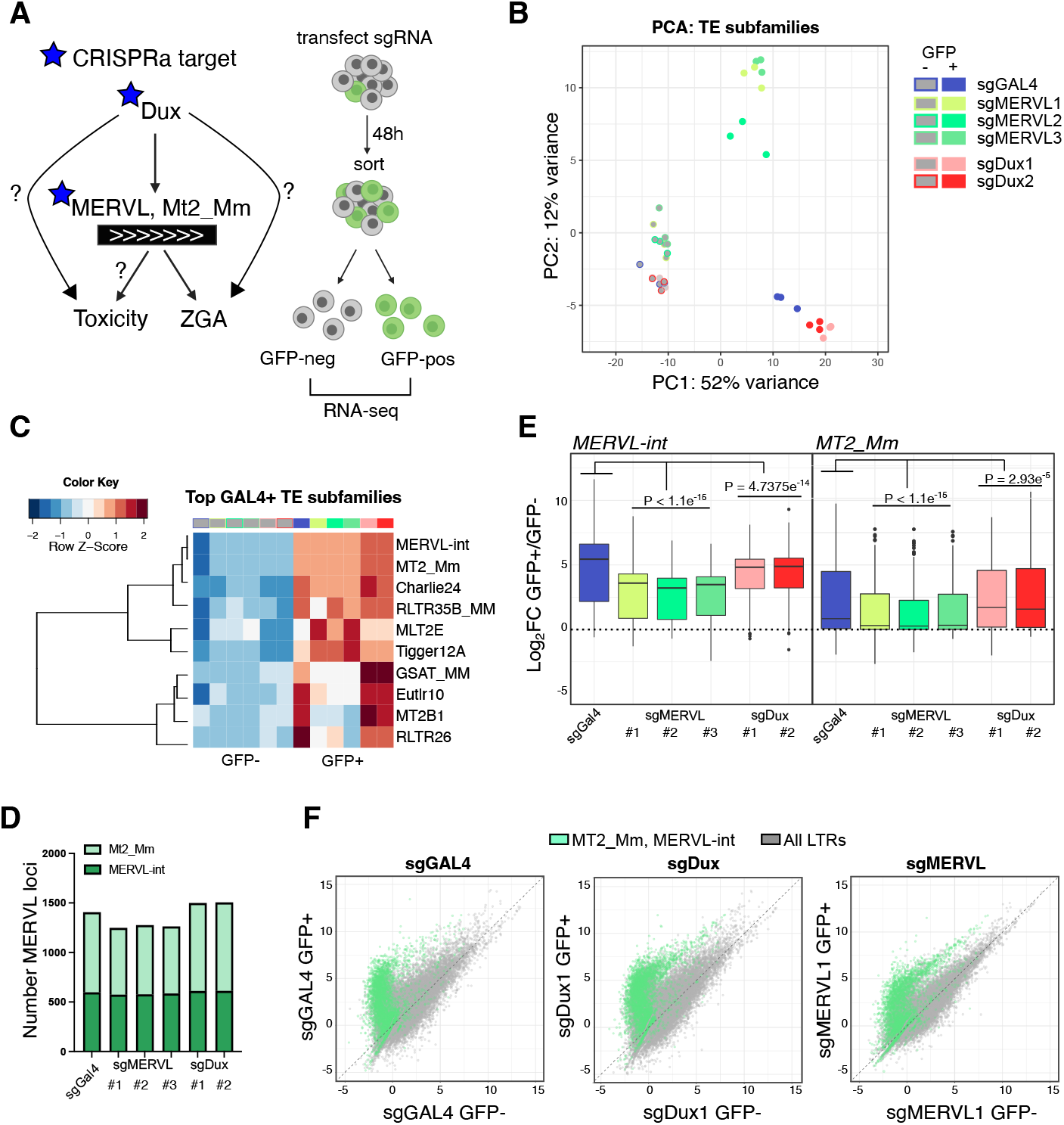
Activation of TEs by Dux- or MERVL-induced 2C-like cells. A) Diagram of CRISPRa experiments and questions. sgRNAs against Dux or MERVL elements were used to probe the relative contribution of distinct portions of this axis to ZGA and to cell toxicity. GFP-positive/negative cells were FACS-purified from sgRNA-transfected CVG ESCs. B) PCA of TE subfamily expression in GFP-negative (-) and GFP-positive (+) cells from the indicated sgRNA transfections. C) Heatmap showing the expression of the top 10 most upregulated TE subfamilies in control 2C-like cells (sgGAL4-GFP+ versus GFP-cells) across all conditions. D) Histogram of the number of significantly-upregulated MT2_Mm and MERVL-int loci in GFP-positive, 2C-like cells compared to their GFP-negative control (FDR < 0.05). E) Boxplot showing log2-fold change (log2FC) in expression of individual loci from MERVL-int or MT2_Mm subfamilies between GFP-positive and GFP-negative samples. P values, Wilcoxon rank-sum test between each condition and sgGAL4, with Bonferroni Correction for multiple comparisons. F) Scatter plot showing normalized expression of all individual LTR loci (grey) and MERVL loci (green) in the indicated conditions.

To further explore TE expression, we investigated individual TE loci (Fig. S2C). As expected, MERVL-int and MT2_Mm loci are induced in all GFP-positive samples, with similar numbers of loci significantly activated in each condition (Fig.2D). While these elements are activated to a similar degree in control versus Dux-induced 2C-like cells, they are consistently induced at slightly lower levels by sgMERVL (Fig.2E-F). The MERVL loci induced by individual MERVL sgRNAs are also broadly overlapping (Fig. S2D). These data confirm that MERVL CRISPRa effectively activates MERVL expression – but that this forms only a portion of total 2C-upregulated repeatome.

### Only a portion of 2C-like genes are MERVL-dependent

The LTRs of TEs such as MERVL can act as powerful promoters of nearby genes, and several 2C-specific genes have been confirmed to be driven by a MERVL MT2_Mm promoter (Macfarlan et al. 2011; Yang et al. 2024). We next therefore examined whether partial TE activation is reflected in the differential activation of genes in CRISPRa-driven 2C-like cells. PCA analysis again revealed that MERVL-induced 2C-like cells possess an intermediate profile between ESCs and control or Dux-driven 2C-like cells (Fig.3A). Of the top 150 upregulated genes in endogenously arising 2C-like cells (GAL4 GFP+), i.e. 2C-like genes, approximately 40% are activated by either MERVL or Dux, while 60% are MERVL-independent (Fig.3B, S3A). In both these cases, Dux-induced 2C-like cells appear, in contrast, very similar to endogenous 2C-like cells. More globally, we found that 558 genes are significantly upregulated in both endogenous and Dux-induced 2C-like cells (“GD”), while 302 genes are also MERVL-activated (“GDM”). 450 genes are also induced only in control 2C-like cells (“G”) (Fig.3C). Very few genes are activated by sgMERVL alone (n=31), revealing that sgMERVL-activated genes form a subset of Dux targets (Fig.3C).

**Figure 3:**
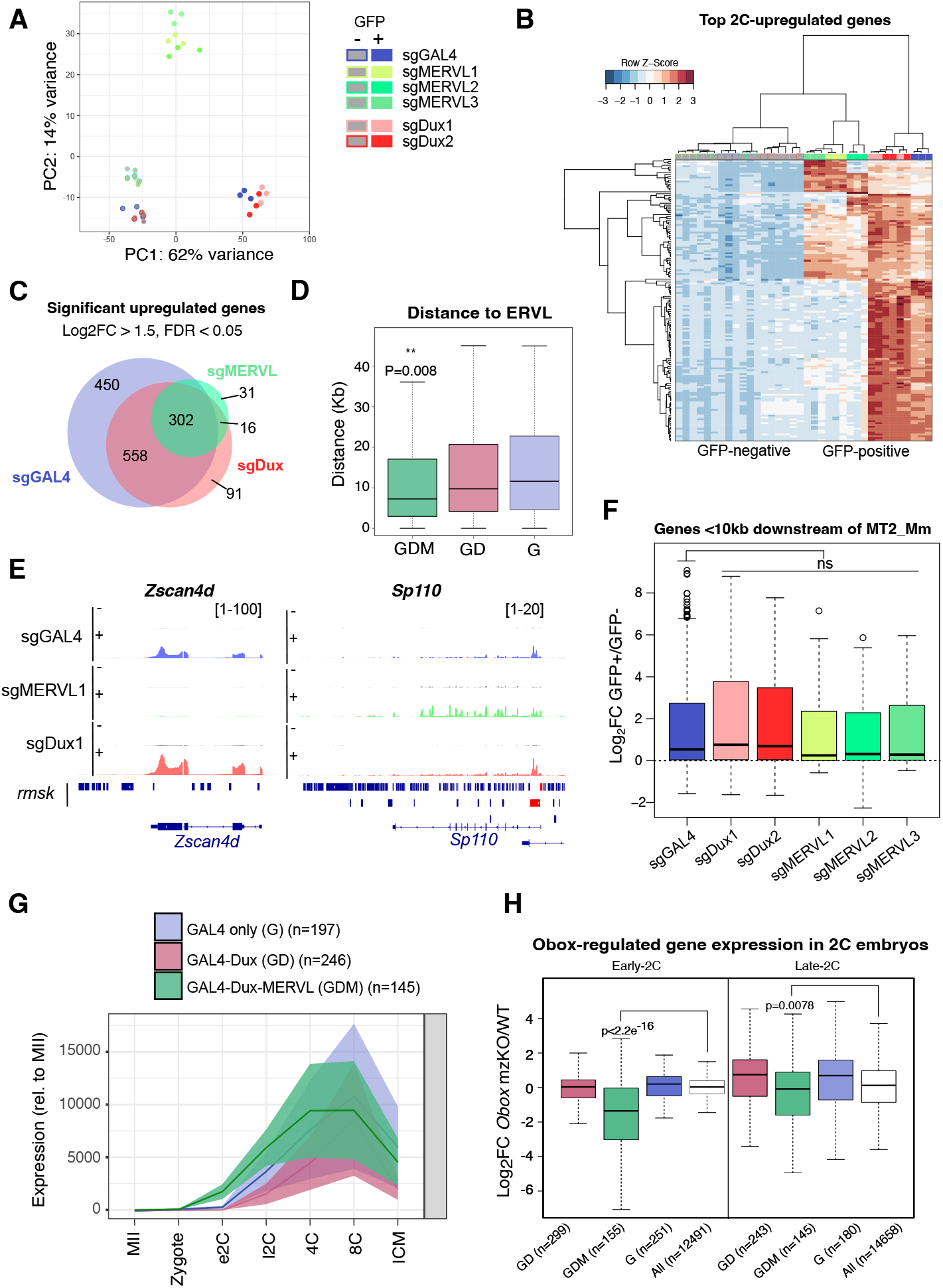
Differential gene induction between MERVL-induced and control 2C-like cells. A)PCA of gene expression in GFP-negative(-) and GFP-positive(+) cells from the indicated sgRNA transfections. B)Heatmap of 2C-like genes (top 200 highest upregulated genes in control (sgGAL4) 2C-like cells) in all samples. C)Venn diagram indicating the overlap in significantly upregulated genes in sgGAL4, sgMERVL, and sgDux GFP-positive over GFP-negative cells (log2FC >1.5, FDR <0.05, in 2 out of 2 or 3 sgRNAs in sgDux or sgMERVL experiments, respectively). D)Boxplot showing absolute distances to the nearest ERVL-family TE from genes in GDM, GD or G gene sets. P <0.008, Wilcoxon rank-sum test between GDM and G. E)Bigwig plots of the indicated genes in representative GFP-positive(+) and GFP-negative(-) sgRNA samples. MERVL elements are highlighted below the tracks in red. F)Boxplots showing log_2_ fold-change in expression of all genes <10Kb downstream from an MT2_Mm across different conditions. Ns = not significant, Wilcoxon rank-sum test with Bonferroni correction. G)Embryo expression dynamics of genes within the indicated gene sets. Lines and shading depict mean +/-s.e.m gene expression, respectively, relative to MII oocyte. Data from (Wu et al. 2016). H)Boxplots showing log_2_ fold-change in expression of the indicated gene sets in *Obox* mzKO 2C embryos over control embryos. Data from (Ji et al. 2023). P values, Wilcoxon rank-sum test between GDM genes and all genes.

A partial induction of the 2C-like transcriptome by MERVL CRISPRa could be due to lower activation of MERVL promoters (Fig.2E), or because many 2C genes are not driven by MERVL. We tested this and investigated the relationship between the MERVL LTR, MT2_Mm, and the gene sets upregulated. We found that GDM genes are significantly closer to an ERVL element than GD or G gene sets (Fig.3D). Moreover, 34.8% of GDM genes are within 10kb of an MT2_Mm, in contrast to only 4.5% or 0.7% of GD or G, respectively (Fig. S3B). For example, the 2C gene, *Zscan4d* is highly upregulated by Dux alone and has no MT2_Mm within 10Kb. Conversely, *Sp110* is most strongly activated by sgMERVL, driven by a MT2_Mm element overlapping its promoter (Fig.3E). Focusing on all genes within 10Kb downstream of an MT2_Mm LTR, these are efficiently induced by direct MERVL activation (Fig.3F, S3C). Taken together, our data reveal that MERVL activation is efficient yet contributes only partially to the 2C transcriptome. In contrast, Dux induces the full spectrum of 2C-like genes, including many MERVL-independent targets.

### MERVL-driven genes are Obox-dependent in ESCs and embryos

MERVL induction is considered a core feature of the 2-cell stage in embryos, and of the 2C-like state in culture. However, our data show that MERVL activation only partially contributes to the genes upregulated in 2C-like cells, unlike Dux activation. Nevertheless, Dux has been found to be ultimately dispensable for embryo development, while MERVL depletion causes cleavage stage arrest (Chen and Zhang 2019; Guo et al. 2019; De Iaco et al. 2020; Sakashita et al. 2023; Yang et al. 2024). These puzzling discrepancies led us to investigate the functional relevance of MERVL-driven (GDM) or MERVL-independent (GD) Dux targets. We first checked expression patterns of these genes in mouse embryos. Post-fertilisation, all 2C-like gene sets exhibit a general upregulation, consistent with a possible role in ZGA or as ZGA targets, with MERVL-dependent genes activated the earliest (Fig.3G). Using data from dual maternal/zygotic knockout (mzKO) of *Dux*, we also asked whether Dux- or MERVL-activated genes are Dux-dependent *in vivo*. All Dux targets showed negligible gene expression changes upon *Dux* mzKO in zygote and 2-cell embryos (Fig. S4A),consistent with the ultimate viability of *Dux*-deleted embryos. These data suggest that other factors can compensate *in vivo* for the absence of Dux at both MERVL-dependent and independent targets.

We hypothesized that MERVL might act as an integrator of many transcription factors, ensuring MERVL-dependent gene activation at ZGA. We investigated Obox factors, which were recently found to be essential activators of MERVL and ZGA in mice. Obox proteins include both maternally and zygotically expressed isoforms, and *Obox* mzKO leads to downregulation of MERVL and ZGA genes and causes early embryonic arrest (Ji et al. 2023; Sakamoto et al. 2024). We first confirmed that *Dux* is not a target of Obox in ESCs or embryos (Fig. S4B). In contrast, *Obox3/5/6* are upregulated in control 2C-like cells and by sgDux, but not by sgMERVL (Fig. S4C). However, these genes are interestingly not affected in *Dux* mzKO embryos (Fig. S4D). Thus, although Dux is sufficient to activate *Obox* isoforms in ESCs, it is not necessary for their expression in embryos.

Next, we investigated whether Obox may collaborate with Dux to activate GDM genes. Using recently-published data (Ji et al. 2023), we found that MERVL-dependent but not MERVL-independent genes are significantly downregulated in both early and late *Obox* mzKO 2C embryos (Fig.3H). Conversely, Obox3/5 overexpression in ESCs directly activates these genes (Fig. S4E). These data demonstrate that MERVL-dependent targets are subject to regulation by Obox factors in addition to Dux. Moreover, their downregulation in *Obox* deficient embryos prior to their arrest could suggest an important role of these genes in ZGA. In contrast, Dux-driven MERVL-independent genes do not change upon *Obox* mzKO. These genes are therefore potentially less important in early embryos. These data further suggest that Dux alone, unlike Obox, is not sufficient to maintain expression of MERVL-dependent genes in 2-cell embryos.

### Dux and MERVL-driven 2C-like cells show different totipotency features

To examine the functional importance of MERVL-dependent and -independent gene expression programmes, we next turned to the known features of 2C-like cells. 2C-like cells have been shown to model several characteristics of 2-cell embryos – including chromatin reorganisation, downregulation of pluripotency factors, and more recently, immature NPB-like nucleoli (Macfarlan et al. 2012; Boskovic et al. 2014; Ishiuchi et al. 2015; Xie et al. 2022). 2C-like ESCs induced from all CRISPRa conditions display downregulation of various pluripotency genes, yet more mildly in sgMERVL cells (Fig.4A). This is also true for Oct4 (*Pou5f1*) protein, with its downregulation most dramatic following Dux induction and only partially downregulated in sgMERVL 2C-like cells (Fig.4B-C, S5A-B). Conversely, nucleoli, which are reprogrammed to resemble NPBs in 2C-like cells (Xie et al. 2022), appear similarly NPB-like in all conditions (Fig.4C-D, S5C). Thirdly, we looked at chromocenter decompaction, which is a marker of the more decondensed chromatin state in 2C-like cells and 2C embryos and accompanied by upregulation of Major Satellite expression (Probst et al. 2010; Ishiuchi et al. 2015; Burton et al. 2020). Strikingly, the expression of Major Satellite RNA, GSAT_MM, is not upregulated by sgMERVL, but robustly activated in control and Dux-driven 2C-like cells (Fig.2C). 2C-like cells driven by sgMERVL indeed show reduced chromocenter decompaction (Fig.4B, S5D). Together, our results suggest that MERVL expression drives only some of the characteristic features of 2C-like cells.

**Figure 4:**
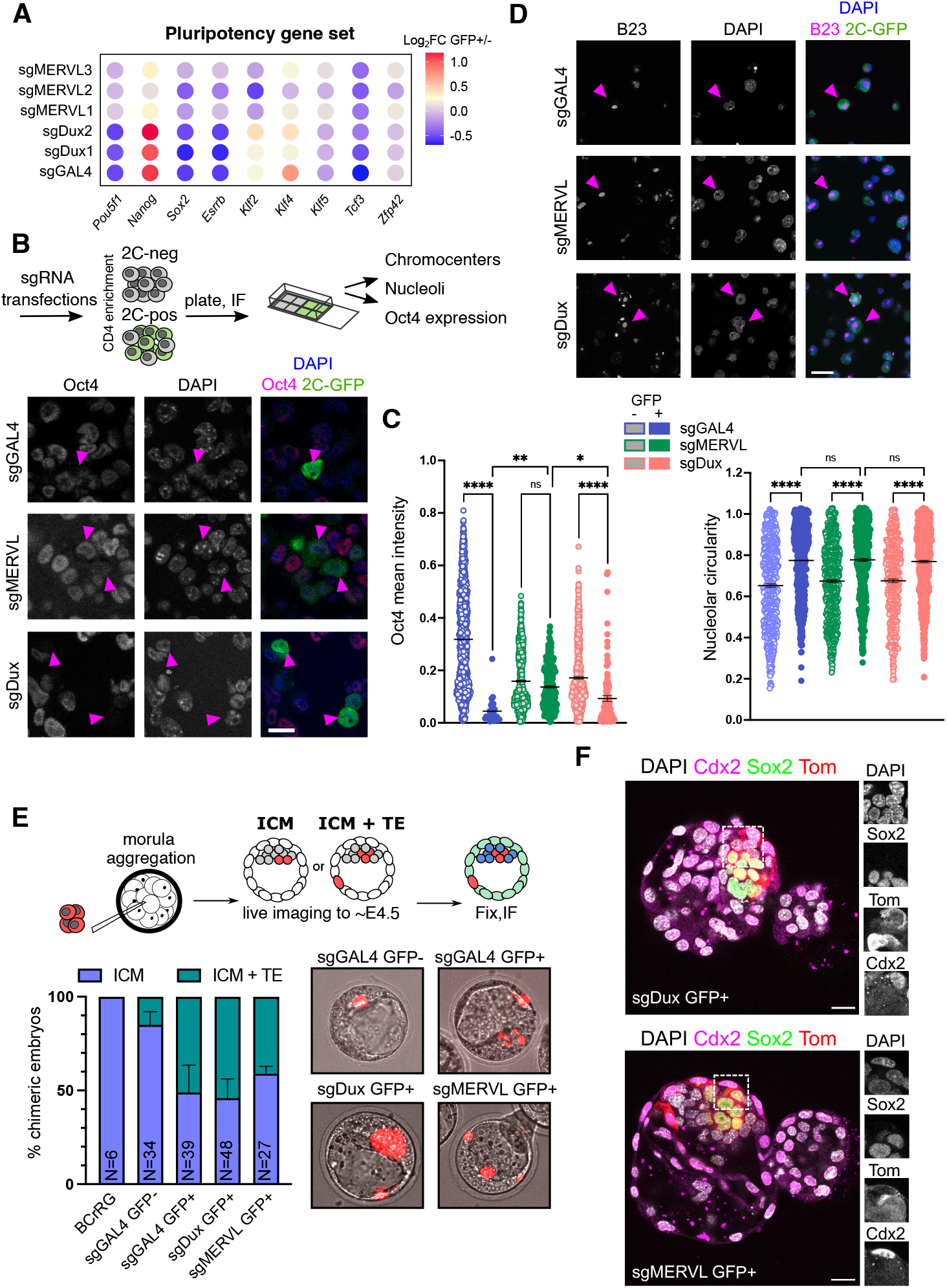
MERVL and Dux-induced 2C-like cells show distinct pluripotency features. A) Bubble plot representing the change in expression of the indicated pluripotency factors in GFP-positive over GFP-negative samples. B) Diagram and representative images of immunofluorescence experiments with sgRNA-induced 2C-like cells. Example 2C-like cells from CD4-based purifications are indicated with pink arrows. Data are representative of at least n=2 experiments. Scale bar, 20 μm C) Quantification of mean Oct4 intensity (left) or nucleolar circularity (right) in ESCs versus 2C-like cells after sgRNA transfection. GFP-negative and positive cells were separated by CD4-based purification and processed for imaging (B,D). Before plotting, data were filtered to exclude outliers based on GFP intensity. P values, one-way ANOVA with Sidak’s multiple comparisons test. Data are representative of 2-3 experiments D) Example images of nucleoli (B23) staining in 2C-like cells from the indicated transfections, as in B. E) Diagram and results of ESC embryo injection experiments. A control ESC line (BCrRG) or TdTomato-labelled CVG ESCs transduced with the indicated sgRNAs were sorted and injected into morula and left to develop to ∼E4.5 with live imaging to follow ESC incorporation. Contribution to ICM or TE was scored by position. Data representative of 3 independent injection experiments (n= embryo number). F) Representative confocal images of equivalent E4.5 chimeras following injection of GFP+ sgMERVL or sgDux transduced cells at the 8-cell stage. Embryos were immunostained for Sox2, Cdx2, and TdTomato (Tom), counterstained with DAPI. Maximum projection of 5 z-sections. Scale bar, 20 µm.

In culture, 2C-like cells are in a totipotent-like state, and unlike ESCs can contribute to both embryonic and extraembryonic lineages when injected into morulae (Macfarlan et al. 2012; Choi et al. 2017; Yang et al. 2020). We next investigated whether there are differences in the expanded fate potential of endogenous, sgDux-or sgMERVL-induced 2C-like cells. CVG ESCs were labelled with tdTomato and transduced with sgRNAs (Fig. S5E) then tdTomato^+^/GFP^+^ cells sorted and injected into WT morulae (Fig.4E). 2C-GFP negative cells, i.e., in an ESC state, were found in the ICM, while control 2C-like cells are present in both the ICM and trophectoderm (Fig.4E), as previously shown (Choi et al. 2017). Interestingly, both sgDux as well as sgMERVL-induced 2C-like cells colonise the ICM and trophectoderm with similar efficiency (Fig.4E). Contribution to both lineages was confirmed with costaining for Sox2 or Cdx2, respectively (Fig.4F, S5F). Therefore, despite the differences in TEs, gene expression and 2C-like characteristics in MERVL-induced 2C-like cells, these cells possess an expanded fate potential that mirrors the 2-cell embryo.

### Dux drives cell death independently of MERVL

While the Dux-MERVL axis promotes totipotency and ZGA, it has also been shown that prolonged or excessive activation can impact cell survival *in vitro* and promote embryo arrest *in vivo* (Percharde et al. 2018; Guo et al. 2019; Olbrich et al. 2021; Xie et al. 2022). Indeed, aberrant DUX4 activation in muscle cells induces an embryonic gene expression programme, ERVL family TE activation, and drives cell death in FSHD (Geng et al. 2012; Shadle et al. 2017). An unanswered question is whether Dux-driven cell toxicity is related to its ability to activate MERVL elements or MERVL-driven genes. Essentially, is MERVL overexpression itself toxic, per se? The downstream mediators of Dux/DUX4-driven cell death are moreover still poorly defined, for example in FSHD. We asked whether FSHD signatures are induced by sgMERVL and examined known FSHD marker gene homologs in our RNA-seq data. We observed that most are specifically upregulated in ESCs by sgDux, but not sgMERVL (Fig.5A). This points to a MERVL-independent network downstream of Dux that is potentially linked to pathology. Tracking CVG ESCs after sgRNA transfection revealed a significant loss of sgDux-transfected CVG ESCs from culture more rapidly than control or sgMERVL-transfected ESCs (Fig.5B). Dux activation is also correlated with a decrease in several cell cycle regulators, which are also known to be affected in FSHD (Fig.5A).

**Figure 5:**
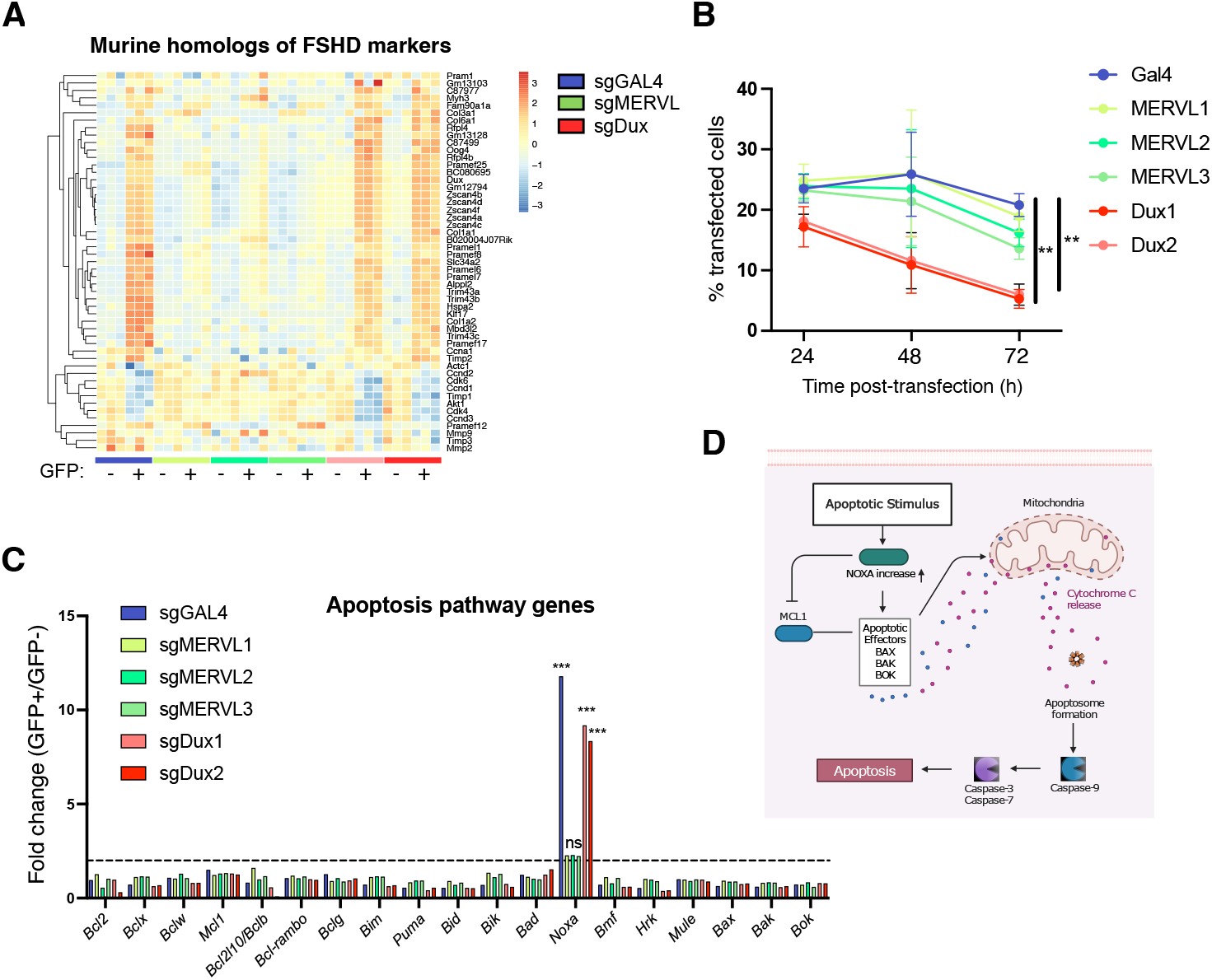
Dux-specific induction of FSHD markers and cell death in ESCs. A)Heatmap of mouse homologs of FSHD marker genes in all GFP-positive and GFP-negative samples. Genes but not samples are grouped by unsupervised hierarchical clustering. B)Histogram showing the percentage transfected (mCherry-positive) cells remaining at the indicated timepoints post transfection. Data are mean +/-s.e.m, n= 3 independent samples from separate experiments. **P < 0.005 at D2 and D3, two-way ANOVA comparing each sample within every timepoint to Gal4 as a control. C)Histogram showing log_2_FC in the indicated apoptosis genes in GFP-positive over GFP-negative samples. *** P < 0.001, Toptable FDR. D)Diagram showing the role of Noxa as a pro-apoptotic sensitizer in cells. Made with Biorender.

We focused on a potential link between Dux and apoptosis and searched for transcriptional alterations in apoptosis-associated effectors (Fig.5C). While the majority of pro- or anti-apoptotic factors show no clear change, the pro-apoptotic sensitizer, *Pmaip1*/*Noxa* (Roufayel et al. 2022), referred to as *Noxa* hereon, is significantly upregulated in Dux-driven 2C-like cells (Fig.5D). Moreover, no significant *Noxa* induction is apparent in 2C-like cells upon MERVL activation. Thus, Dux activation leads to reduced cell numbers and changes in the apoptosis pathway that are independent of MERVL activation and associated with poor survival of these cells.

### Noxa contributes to Dux-driven apoptosis in ESCs

The induction of *Noxa* suggested it may be a potential mediator of Dux-driven apoptosis in ESCs. To test this, we first established a Tet-ON system to allow for inducible Dux expression (iDux) in an independent 2C-GFP line (Percharde et al. 2018) (Fig.6A). Doxycycline (Dox)-mediated Dux induction leads to robust induction of 2C-like cells by 24h, as expected (Fig.6B, S6A), and these cells are lost from ESC culture in a similar manner to Dux CRISPRa ESCs (Fig.6C). *Noxa* is upregulated upon Dux induction (Fig.6D), and ChIP-seq analysis revealed that Dux binds at two *Dux* motifs upstream of the *Noxa* promoter (Eidahl et al. 2016) (Fig.6E). In contrast, no MT2_Mm is found within 50kb of *Noxa*. These data reveal that the apoptotic factor *Noxa* is a direct Dux target in ESCs.

**Figure 6:**
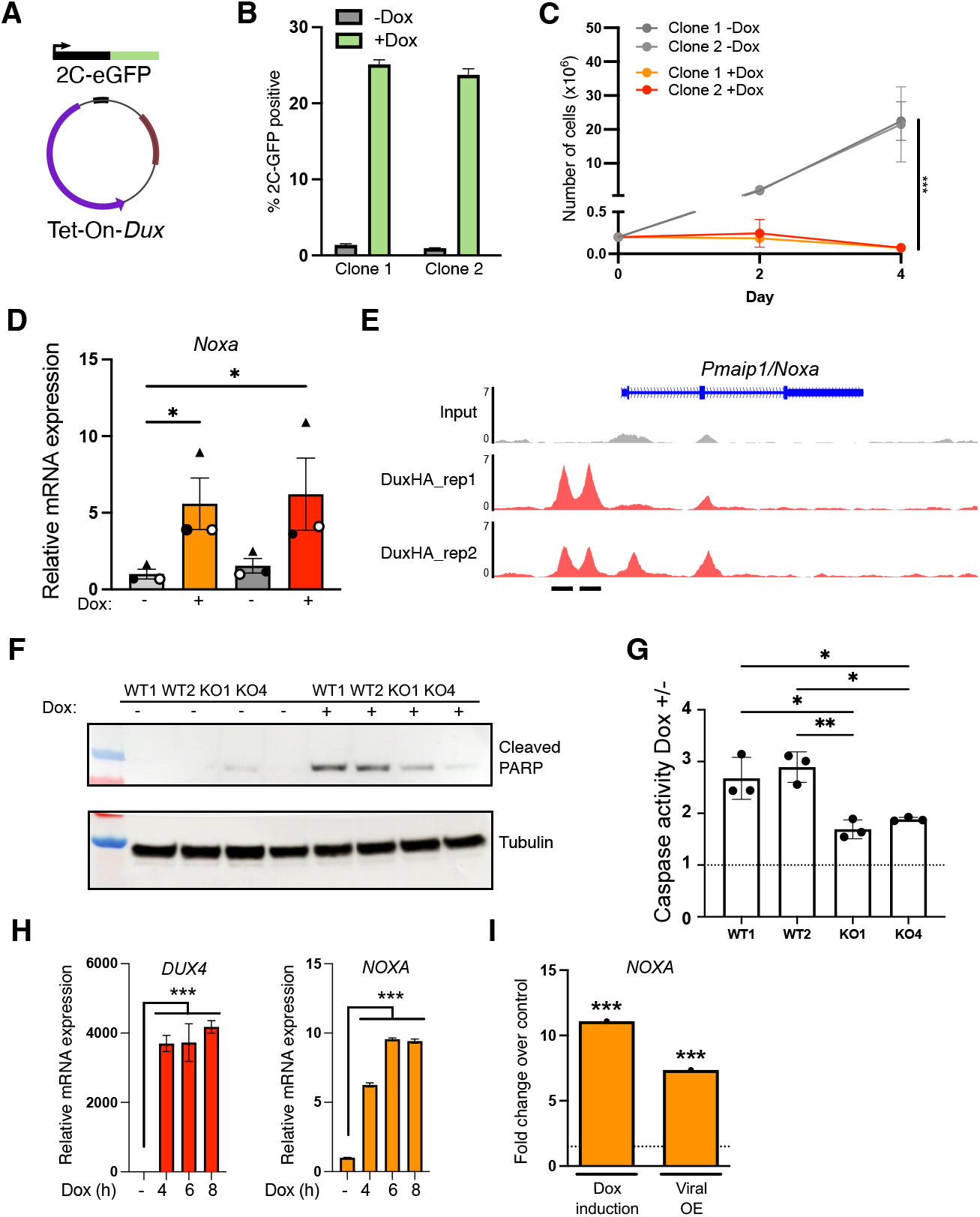
Direct Dux activation of pro-apoptotic Noxa/NOXA. A) Diagram showing Dox-inducible-Dux plasmid, introduced by lentiviral transduction into an independent 2C-GFP reporter ESC line (Percharde et al. 2018). Made with Biorender. B) Histogram showing the percentage of 2C-GFP positive cells quantified by flow cytometry following Dox-mediated Dux induction. Data are mean +/-stdev, n=3 independent experiments on two different clones. C) Graph of ESC proliferation in cells cultured with or without Dox in order to induce Dux expression. Data are mean +/-stdev, n=3 independent experiments, ***P < 0.001, 2-tailed student’s t test. D) qRT-PCR analysis of *Noxa* in ESCs following Dux induction. Data are mean +/-n=3 independent experiments, *P<0.05 two-way ANOVA. E) ChIP-seq analysis of the *Noxa/Pmaip1* locus, showing peaks of Dux binding at two *Dux* motifs (Eidahl et al. 2016). ChIP-seq data from (Hendrickson et al. 2017). F) Western blot of WT or *Noxa* KO ESCs 48h following Dux induction. 2 independent clones are shown, representative of at least n=2 experiments. G) Caspase 3/7 assay on *Noxa* WT or KO ESCs 24 h after Dox-induction of Dux. Data are shown as fold increase in activity of Dox-treated cells over untreated cells, n=3 independent experiments. *P <0.05, **P<0.005 one-way ANOVA. H) *DUX4* and *NOXA* expression in a Dox-inducible DUX4 human ESC line (Yoshihara et al. 2022). Data are mean +/-s.e.m, n=3 wells, representative of 2 experiments. P values are from a one-way ANOVA with Dunnett’s multiple comparisons test. I) Expression analysis of *NOXA* upon DUX4 induction in two pre-clinical myoblast models of FSHD. ***P <0.001, Toptable analysis, FDR <0.05. Data analysed from (Jagannathan et al. 2016).

Noxa is a pro-apoptotic sensitizer, and it has been shown that altering the balance between levels of pro and anti-apoptotic factors can push cells into apoptosis (Fig.5D). To test whether Noxa might contribute to Dux-induced apoptosis, we generated *Noxa* KO clones in iDux 2C-GFP ESCs (Fig. S6B-C). We examined markers of apoptosis upon Dux induction in WT and KO clones, revealing that levels of cleaved-PARP are lower in the absence of Noxa (Fig.6F). This is supported by reduced Caspase 3/7 activity in *Noxa* KO ESCs (Fig.6G). These data suggest that Noxa is a Dux-induced mediator of apoptosis and may be partly responsible for the poor survival of Dux-induced 2C-like cells.

Finally, we investigated if the Dux-Noxa relationship is conserved in human cells. We took advantage of recently described iDUX4 human ESCs (hESCs), where the Dux homolog, DUX4, is induced upon Doxycycline (Dox) addition (Yoshihara et al. 2022). A short pulse of Dox efficiently activates DUX4 target genes that are detectable 12 hours following Dux induction; however, no upregulation of *NOXA* was seen (Fig. S6D). Interestingly, these hESCs also do not appear to undergo extensive cell death (Fig. S6E). We tested if sustained DUX4 expression might instead induce *NOXA* and found that both *DUX4* and *NOXA* expression is significantly upregulated by 4 hours following continuous Dox-induction, with *NOXA* levels peaking at 12 hours (Fig.6H, S6F). In agreement, these hESCs exhibit signs of cell death by 12-14 hours of Dox treatment (Fig. S6E). Interestingly, *NOXA* is also significantly upregulated in two pre-clinical myoblast models of FSHD (Fig.6I). Taken together, our results reveal that Dux alone induces cell death in ESCs independently of MERVL, and suggests that Dux/DUX4-driven activation of NOXA may be a conserved axis contributing to apoptosis in mouse and human cells.

## Discussion

The transient upregulation of MERVL transposons is a known, striking feature of murine ZGA. It also typifies the 2C-like cells *in vitro* that have been found to resemble the 2-cell embryo. Despite this, the mechanisms and functions of MERVL activation are still poorly understood. The relationship between MERVL and its transcriptional activator, Dux, is also a puzzle. Dux expression is both sufficient and necessary for 2C-like cell induction and spontaneous appearance within ESC culture (Hendrickson et al. 2017; De Iaco et al. 2020). However, multiple groups have demonstrated that *Dux* mzKO embryos are ultimately viable, with sustained MERVL and 2C gene expression (Chen and Zhang 2019; Guo et al. 2019; De Iaco et al. 2020; Bosnakovski et al. 2021). It was thus unclear which parts of this network are the most important for 2C-like features and for development. Conversely, it was also unclear whether the developmental arrest or apoptosis seen upon Dux overexpression in various contexts (Percharde et al. 2018; Guo et al. 2019; Xie et al. 2022) is linked to MERVL, or due to independent functions of this factor.

Our study sheds light on the Dux-MERVL relationship and its relation to 2C-like characteristics and Dux-induced toxicity. Using a CRISPRa system, we have compared and contrasted the functions of these factors. First, we find that MERVL CRISPRa efficiently activates the majority of functional MERVL promoters, and yet this comprises only a portion of the endogenous 2C-like transcriptome and repeatome. In contrast, Dux-driven 2C-like cells are remarkably similar to endogenously-occurring 2C-like cells in both their TE and gene expression. Our results reveal that the basis of these differences is MERVL-independent; genes promoted by MERVL are efficiently activated under all experimental conditions. Despite the high number of genes and TEs only upregulated in control and Dux-induced 2C-like cells, we suggest that these less important in ZGA. Such MERVL-independent genes are not affected upon *Obox* deletion, which induces early embryo arrest and defective ZGA (Ji et al. 2023). In contrast, MERVL-dependent 2C genes are significantly downregulated in *Obox* mzKO, consistent with a role for these genes in the early embryo. Taken together, our data define a minimally required set of 2C genes, driven by MERVL elements. We suggest that a key function of the MT2_Mm promoter is as an integrator of multiple transcription factors (eg Obox and Dux), providing several ways of inducing these 2C genes for ZGA.

At a molecular level, MERVL-induced 2C-like cells display similar immature nucleolar morphology as endogenous 2C-like cells. Interestingly however, they only partially downregulate Oct4 and fail to decondense chromocenters or upregulate Major Satellite RNA, suggesting only a partial reprogramming of 2C-like chromatin. Nevertheless, both MERVL- and Dux-induced 2C-like cells can contribute to ICM and trophectoderm when injected into morulae – a key functional characteristic of the 2C-like state that mimics totipotency in the 2-cell embryo. Future experiments may shed light into the mechanisms and functions of Dux-driven chromatin reprogramming in these distinct scenarios.

Our results build upon previous studies that have identified factors that can induce 2C-like and totipotency characteristics (Choi et al. 2017; Hu et al. 2020; Yang et al. 2020), either upstream or downstream of Dux (Sugiyama et al. 2024). Here, through direct comparison we reveal that direct MERVL or Dux upregulation confers distinct features of totipotency. Moreover, by defining genes and repeats specific to each part of the axis, our data point to a minimal set of MERVL targets responsible for these totipotency characteristics. Investigating these factors in future may reveal novel proteins important in development for essential processes such trophectoderm specification and ZGA.

This work, as well as others, points to some interesting differences between ESC models of the 2-cell stage and 2-cell embryos. Although Dux potently activates both MERVL-dependent and independent genes in 2C-like cells, these are not Dux-dependent in embryos. The reports of viable *Dux* KO mice have led to suggestions that studying this factor – or 2C-like cells – may have limited interest or utility in development. However, we argue that this dichotomy serves to highlight the multi-pronged mechanisms in place to ensure ZGA occurs. A system where multiple factors all activate the same core genes provides the opportunity for redundancy and robustness in an essential developmental process. Indeed, newly identified ZGA factor Nr5a2 activates itself as well as other nuclear receptors such as Esrrb that bind the same motif, upregulating 75% of ZGA genes (Gassler et al. 2022). Multiple Obox factors are expressed in 1-2 cell embryos, and Obox4 compensates for the loss of Dux (Guo et al. 2024). This raises the question of whether Obox and Dux act truly redundantly or if additional factors present in embryos, but missing from embryonic stem cells (ESCs), further promote MERVL gene expression. Differences in both chromatin organisation as well as factor availability may underlie contrasting dependencies in 2C-like cells versus 2-cell embryos. For example, both Nr5a2 and Esrrb bind distinct sets of enhancers in 2-cell embryos and ESCs (Gassler et al. 2022). Investigating these discrepancies will shed further light on the mechanisms of ZGA and early development.

The flip side to the Dux-MERVL axis is that its activation is only tolerable for a very short period. Ectopic *Dux* overexpression halts embryo development (Guo et al. 2019), while preventing Dux repression by various means is sufficient to cause 2-cell arrest (Percharde et al. 2018; Xie et al. 2022; Vega-Sendino et al. 2024). In ESCs, Dux also induces DNA damage at CTCF motifs (Olbrich et al. 2021). However, the role of MERVL in Dux-dependent cell death was previously unknown. Here, we demonstrate that Dux-dependent cell death is MERVL-independent. Additionally, we identify the apoptotic sensitizer, Noxa, as a direct Dux-target. *Noxa* deletion reduces Dux-dependent apoptosis in ESCs, and we show that DUX4 also induces *NOXA* in hESCs. Moreover, *NOXA* is upregulated in pre-clinical models of FSHD. These data suggest NOXA may be a promising therapeutic target to reduce myoblast death and potentially, muscle loss in FSHD. This could function either as a standalone treatment or in conjunction with future therapies that could directly repress DUX4 expression (Le Gall et al. 2020) or those that might inhibit DUX4 signalling but that have not themselves reduced cell death (Oliva et al. 2019).

Given the above, an intriguing question is how Dux/DUX4 expression is tolerated in normal development in both mice and humans. We hypothesize that this is because its expression is only transient in these biological contexts. The 2C-like state is also temporary in cell culture, with endogenously arising 2C-like cells soon transitioning back into ESCs (Macfarlan et al. 2012; Eckersley-Maslin et al. 2016). Thus, even though Noxa is upregulated in 2C-like cells (Fig.5C), this may be brief enough to avoid triggering cell death.

Indeed in human ESCs, *NOXA* levels only rise after sustained DUX4 activation, unlike other targets such as *ZSCAN4* that are detected after a brief pulse of DUX4. The induction of Noxa/NOXA and so apoptosis may serve as a natural mechanism to prevent prolonged activation of these pathways. Learning more about the processes responsible for totipotency and ZGA, as well as those that trigger pathology, will enhance our understanding of healthy development.

## Materials and Methods

### Cell culture

Mouse embryonic stem cells were cultured on 0.2% gelatine-coated (G1393-100ML, Sigma-Aldrich) dishes in DMEM + Glutamax (31966047, Gibco) supplemented with 10% FBS (A5256801, Gibco), 2-mercaptoethanol (31350010, Gibco), non-essential amino acids (11140050, Gibco) and LIF (ESG1107, Millipore). Upon establishment, CVG ESCs were cultured in the presence of 10 µg/ml Blasticidin (11583677, ThermoFisher) and 250 µg/ml G418 (11811023, ThermoFisher). For cells inducible by doxycycline, a final concentration of 1 µg/ml of doxycycline was used (A4052-APE-1g, Stratech Scientific). Length of induction varied between experiments and is indicated in the respective figure legend.

Doxycycline-inducible DUX4-TetOn hESCs (iDUX4 hESCs) were previously generated from the H9 (WA09) human embryonic stem cell line (Yoshihara et al. 2022). Cells were cultured on Geltrex (A1413302, Life Technologies) coated 6-well tissue culture dishes in Essential 8 medium (A1517001, Thermo Fisher Scientific) at 37°C with 5% CO2, and passaged every 3–5 days after approximately 5 minutes of incubation with 0.5 mM EDTA (15575020, Life Technologies).

To induce DUX4 expression, DUX4-TetOn hESCs were treated with 1 μg/mL doxycycline in Essential 8 medium (5% CO2, 37°C). For pulsed induction, cells were incubated with doxycycline (1 μg/mL) for 15 minutes, washed three times with Essential 8 medium, and then incubated in fresh Essential 8 medium for 12 hours. For continuous induction, cells were exposed to doxycycline (1 μg/mL) in Essential 8 medium (5% CO2, 37°C) for varying durations (0, 4, 6, 8, or 12 hours) before being harvested.

### Cell line generation

CRISPRa activity in E14 ESCs was achieved by safe-harbor integration of the dCas9-VPR transgene (Chavez et al. 2015) driven by a CAG promoter and linked to Blasticidin resistance into the *ROSA26* locus. The dCas9-VPR knock-in vector containing ∼500 bp 5’-, and 3’-homology arms was linearized by Ahd I digestion and recovered following phenol/chloroform extraction and ethanol precipitation. 10 µg of linearized targeting vector + 5 µg of pX458-Cas9-ROSA sgRNA was introduced into ESCs by liposome-based transfection with JetPrime (Polyplus-transfection SA). Following transfection, ESCs were plated on gelatine-coated 10 cm plates at low density to ensure clonal expansion. 48 hours after plating the media was refreshed to ESC media plus 2.5 μg/mL Blasticidin to initiate selection for recombinants. Following 8-10 days of Blasticidin selection, individual resistant colonies were isolated and transferred to individual wells of a gelatinized 96-well plate for further expansion and PCR screening. Individual clones were further screened for homologous recombination by PCR genotyping of gDNA across the ROSA 5’-homology arms. Heterozygous recombinants were subsequently screened for transcriptional activation activity by qRT-PCR following lentiviral delivery of validated sgRNAs against *Hbb-bh1* and *Ttn* (Chavez et al. 2016).

To incorporate the 2C-GFP/CD4 reporter into these cells, a single heterozygous *Rosa26*^dCas9-VPR/+^ clone was targeted with our previously-described 2C-GFP-t2a-mCD4 reporter construct (Xie et al. 2022) by electroporation, as previously described. 50 µg of each reporter was linearized with DraIII digestion and cleaned up by phenol-chloroform extraction followed by ethanol precipitation prior to electroporation. Electroporated cells were plated at low density to ensure clonal density would be preserved. Electroporated cells were subjected to geneticin/G418 selection (250 µg/mL) 24 hours after electroporation. Selection was maintained 7-8 days until individual resistant colonies emerged. A single positive clone with expected 2C-GFP behaviour was used for all subsequent experiments (CVG ESC line)

To generate Dox-inducible Dux overexpressing mouse ESCs with a 2C-GFP reporter, lentivirus containing Doxycycline-inducible plasmid construct (pCW57.1-mDux-CA, a gift from Stephen Tapscott (Addgene plasmid #99284; http://n2t.net/addgene:99284; RRID:Addgene_99284) was produced in 293T cells and used to infect an independent 2C-GFP reporter ESC line (Percharde et al. 2018). Infected cells were selected with 1 µg/mL puromycin. Individual clones were picked, expanded and their response to doxycycline induction was tested. Two of the most responsive clones were selected and clonally expanded.

In order to knock out *Pmaip1/Noxa*, gRNAs targeting both ends of the Pmaip1 gene were cloned into the pX458 plasmid carrying either a BFP or GFP reporter and transfected into Dux-inducible 2C reporter cells (Table S1). Double-positive cells were then sorted and individual clones were picked and screened by PCR for the *Pmaip1* gene deletion.

### CRISPRa gRNA design and cloning

The locations of MT2_Mm family members were extracted from the UCSC repeatmasker track for mm10 and the corresponding nucleotide sequences extracted from the genome sequence. Candidate SpCas9 sgRNAs, using an NGG PAM sequence, were identified for these elements using FlashFry (McKenna and Shendure 2018), allowing for no mismatches. The MT2_Mm consensus sequence was downloaded from Dfam (DF000004155) (Storer et al. 2021) and, finally, guides with highest coverage in the genome and with exact matches to the consensus sequence were shortlisted.

Dux sgRNAs were designed using the Benchling CRISPR design tool, inputting the promoter sequence and CDS start of *Dux (ENSG00000058537)* as input. The two sgRNAs giving the highest Dux induction in transfection experiments were selected for all experiments in this study. For all transfection and transduction experiments, sgRNAs cloned into Aar I sites (ThermoFisher) downstream of a U6 promoter in the lentiviral expression vector mp783 were used, which also contains a separate EF1a-driven PuroR-t2a-mCherry reporter cassette.

### Transfection and lentivirus transduction

Transfections were done using Lipofectamine 2000 (11668027. Invitrogen) in Opti-MEM (31985062, ThermoFisher) according to manufacturer’s instructions. Cells were trypsinized and collected 24 h or 48 h post-transfection depending on experimental design.

For lentivirus generation, Lenti-X 293Ts (632180, Takara) were maintained in high glucose DMEM Glutamax (31966047, Gibco), 10% FBS (A5256801, Gibco), 1X MEM non-essential amino acids (11140050, Gibco) and 0.1 mM 2-Mercaptoethanol (31350010, Millipore). The day before transfection, Lenti-X 293Ts were seeded on 0.1% (w/v) Poly-L-lysine coated (P8920, Sigma) dishes at a density of 105,000 cells cm^2^ and allowed to adhere overnight. sgRNA-containing plasmids were transfected using Lipofectamine3000 (L3000008, ThermoFisher) with packaging plasmids psPAX2 and pMD2.G. 6 h post-transfection, plates were refreshed with lentivirus packaging media (high glucose DMEM Glutamax, 5% HI-FBS, 1X MEM non-essential amino acids and 0.1 mM 2-Mercaptoethanol). 48 h post-transfection, the media was collected, and centrifuged to remove debris. The supernatant was then passed through a 0.45 μm filter and stored at 4°C. Media was refreshed for the cells and 24 h later, the media was again harvested as above. Lentiviruses collected on 48 h and 72 h timepoints were concentrated using the 100kDa Amicon ultracentrifugal units (UFC810024, Millipore), spun at 1500g, maintained at 4°C. Concentrated lentiviruses were aliquoted and snap frozen. Once thawed, unused portions of the aliquots were discarded.

For lentivirus transduction, 10^5^ cells were plated in a 6-well plate coated with 0.2% gelatine. The next day, culture media was changed and 8 µg/mL polybrene (7711, Tocris) was added to the media prior to the addition of 15 µl of concentrated virus. Media was changed every day until cells were trypsinized and collected (48 h or 72 h post-infection depending on experimental design).

### Flow cytometry

Cells were trypsinized and washed with PBS before being resuspended in FACS buffer (PBS, 3% FBS, 1mM EDTA) and passed through a 35 µm strainer cap. Single-cell suspensions were evaluated on an LSRII Flow Cytometer System (FACSymphony A3 - BD Biosciences) equipped with FACSDiva software.

### Preparation of 2C- and 2C+ cells for RNA-seq

CVG ESCs were transfected with the different gRNA plasmids using Lipofectamine 2000 (11668027, Invitrogen) according to manufacturer’s instructions. Culture media was changed the next day. Cells were trypsinized 48 h post-transfection and 2C- and 2C+ cells (GFP- and GFP+ respectively) were isolated by FACS. Cells were then pelleted and RNA purified using the RNeasy Mini Kit (74104, Qiagen). RNA was sent to Novogene for RNA-seq library preparation and sequencing.

### RT-qPCR

RNA was isolated using the RNeasy Mini Kit (74104, Qiagen) according to manufacturer’s instructions. Up to 1 µg of RNA was used for cDNA synthesis using the High-Capacity RNA-to-CDNA kit (4387406, ThermoFisher). qPCR was done using KAPA SYBR FAST Rox Low (KK4622, Merck) on a QuantStudio5 (Applied Biosystems). Quantification of gene expression was achieved by normalising to *Rpl7*. All primers used for qPCR can be found in Table S1.

### RNA-sequencing

RNA was purified from cells harvested in 3 independent CRISPRa and FACS experiments and verified by Bioanalyser analysis to have a RIN of >7.0. RNA-seq libraries were prepared by Novogene and sequenced PE 150bp, with an average of at least 30M reads per sample. Reads were processed using cutadapt v4.7 (Martin 2011) to remove Illumina adapters, trim, and filter on quality with the settings: -a AGATCGGAAGAGCACACGTCTGAACTC CAGTCA, -A AGATCGGAAGAGCGTCGTGTAGGGAAAGAGTGT, -n 1, -m 31, --nextseq-trim 20, -- max-n 0. Processed reads were aligned with HISAT2 v2.2.1 (Kim et al. 2019) to GRCm39 with settings adjusted to report up to 100 multiple alignments for each read pair: --rna-strandness RF --no-discordant -- no-mixed --no-unal -k 100 --score-min L,0,- 0.66 --pen-noncansplice 20 --dta.

Gene and TE locus-level counts were obtained with TElocal (TEtranscripts (Jin et al. 2015)) using settings to allow distribution of multimapping reads using maximum likelihood estimation: --stranded reverse -- mode multi. GRCm39.110 (Ensembl) annotations were used for genes. TE annotations were produced using RepeatMasker v4.1.5 configured with HMMER v3.3.2 (Wheeler and Eddy 2013) to scan the GRCm39 genome using Dfam3.7 (Storer et al. 2021) HMM TE models in sensitive mode: -s -no_is -norna - spec “mus musculus”.

Within R v4.3.3, gene count data was used assess inter-sample relationships using hierarchal clustering and PCA. Additionally, the impact of technical experimental factors was assessed using variance partitioning (variancePartition v1.30.2 (Hoffman and Schadt 2016)). Following these assessments, the RUVs method (RUVseq, (Risso et al. 2014)) was used to remove the replicate batch effect by subsequently integrating the returned correction factors into the statistical model (Table S2).

Gene and TE analysis were performed using DESeq2 v1.40.2 (Love et al. 2014). TE subfamily analysis was performed by summing the counts from individual TE loci. Within DESeq2, independent hypothesis weighting was conducted to optimise the power of *p value* filtering using IHW v1.28.0 (Ignatiadis et al. 2016) and log2 fold-change shrinkage was performed using ashr v2.2-63 (Stephens 2017). Multiple correction testing adjustments were optimised by passing FDR and log2FC thresholds at the point of results generation.

### MACS-Immunofluorescence

MACS cell separation was performed using the EasySep™ Mouse CD4 Positive Selection Kit II (18952A, StemCell Technologies). Briefly, CVG ESCs were trypsinized, washed once with PBS and resuspended in PBS FACS buffer (PBS, 3% FBS, 1 mM EDTA) at a concentration of 10^8^ cells per mL. 2C negative (flowthrough) and 2C positive (beads) cells were purified according to manufacturer’s protocol and plated on matrigel-coated chambered slides (354108, Falcon) for immunofluorescence experiments. 1 h after plating, cells were fixed for 10 minutes using 4% paraformaldehyde in PBS. For immunofluorescence, cells were washed with PBS and then blocked and permeabilized using a blocking buffer (PBS, 10% Donkey serum and 2.5% BSA) supplemented with 0.4% Triton-X for 30 minutes at room temperature. Primary antibodies in blocking buffer were added for 1h at room temperature or overnight at 4°C (Table S1). After the incubation with the primary antibodies, cells were washed 3 times with PBS and then incubated with the corresponding secondary antibodies for 1 h at room temperature. Cells were finally washed 3 times with PBS, before the slides were covered with Vectorshield plus DAPI, mounted and sealed. For graphs in Fig.4C, to correct for low enrichment following MACS, outliers were removed based on unexpected GFP level (eg GFP-negative cells removed from the 2C-positive condition). See also Fig. S5A (unfiltered Oct4 data).

### Western Blot

Cells were lysed in RIPA buffer (150 mM NaCl, 5 mM EDTA, 50 mM Tris ph8.0, 1% NP-40, 0.5% sodium deoxycholate, 0.1% SDS). Protease inhibitors (78429, Thermo Fisher Scientific) were added freshly before every use. After the addition of 4X Bolt LDS

Sample Buffer (B0007, Invitrogen), samples were heated for 5 min at 95°C and loaded on a 4-12% Tris-Glycine gel (XP04122BOX, Invitrogen). The transfer onto a PVDF membrane (IB24001, Invitrogen) was done using the iBlot 2 (Invitrogen) following manufacturer’s instructions. Membranes were blocked in 5% non-fat milk (42590.02, Serva) in TBST for 1 hour and incubated with primary antibodies (Table S1) overnight at 4°C. Blots were washes three times with TBST and incubated with the corresponding HRP-conjugated secondary antibodies (Table S1) for 1 hour at room temperature in TBST. Blots washed three times in TBST and imaged using the ImageQuant 800 (Amersham) following manufacturer’s instructions.

### Embryo micromanipulation

Animal studies were authorized by a UK Home Office Project Licence under the Home Office’s Animals Act 1986 and carried out in a Home Office designated facility. B6CBA-F1 females (6-9 weeks old) were superovulated with 5 IU PMSG intraperitoneal injection followed by 5 IU hCG intraperitoneal injection 48 h after. Pre-implantation embryos were collected at embryonic day 2.5 (E2.5) and cultured in KSOM medium (MR-101-D, Sigma) at 37°C and 5% CO2 until micromanipulation. For micromanipulation, E2.5 mouse embryos were placed in pre-warmed M2 medium (M7167, Sigma) under light mineral oil (ES-005-C, Sigma). Embryo cell injections were performed on an inverted microscope (Leica DMi8) using Narishige micromanipulators and hydraulic microinjectors. Eppendorf capillaries were used during the micromanipulator for embryo holding (VacuTip I, 5195000036, Eppendorf) and for cell injection (TransferTip (ES), 5195000079, Eppendorf). Five to six ESCs were injected per E2.5 mouse embryo. After microinjection, embryos were incubated in KSOM medium (MR-101-D, Sigma) at 37°C and 5% CO2. A control ESC line constitutively expression tdTomato and OCT4-GFP (labelled as BCrRG) shown to give good ICM contribution (Sepulveda-Rincon et al. 2024) was used as a control.

### Live-cell imaging of pre-implantation chimeras

Time-lapse imaging was performed on an Olympus IX83 microscope equipped with a Hamamatsu ORCA-Flash 4.0 camera and environmental chamber kept at 37°C with a 5% CO2 supply. Z-stacks were collected with a step-size of 5 μm, every 2 h for 48 h using a UCPlanFN N 20x/0.70 objective for embryos. Embryos were placed into micro-insert 4-well fultrac dish (80406 or 80486, Ibidi) in KSOM medium (MR-101-D, Sigma). Embryos were scored at blastocyst stage immediately before moment of hatching for contribution of TdTomato+ ESCs to either ICM and/or TE, based on position.

### Immunofluorescence imaging of pre-implantation chimeras

Embryos were fixed in 4% PFA in PBS for 20 min at room temperature, followed by three five-minute washes in PBS containing 0.1% PVA (PBS-PVA). Samples were permeabilized for 30 min in 0.5% Triton X-100 in PBS at room temperature, then blocked for 1.5 h at room temperature in 5% BSA-PBS. Samples were incubated with primary antibodies (1:100 in 5% BSA-PBS, Table S1) in 10 µL drops in a humidified Terasaki plate (Greiner Bio-One) overnight at 4°C. Embryos were washed three times in PBS containing 0.1% Tween-20 and 0.1% PVA (PBST-PVA) for 5 min. Samples were incubated in secondary antibodies (1:500 in 5% BSA-PBS) (Table S1) in 10 µL drops in a humidified Terasaki plate for 1 h in the dark. Samples were washed three times in PBST-PVA for 5 min and mounted in Vectashield with DAPI (H-1200-10, Vector Laboratories) under #1 coverslips supported by Dow Corning high-vacuum silicone grease (Z273554-1EA, Sigma-Aldrich) on glass slides. Confocal imaging was performed on a Leica Stellaris microscope with a 40x glycerol objective and acquired in 1 µm Z-stacks.

### Caspase 3/7 Assay

10^4^ ESCs per condition were seeded on 0.2% gelatine-coated 96-well plates. 24 h later, 1 μg/mL doxycycline was added for an additional 24 h to induce Dux expression. The next day, the caspase 3/7 assay was done using the Caspase-Glo 3/7 Assay System (Promega, G8092) according to manufacturer’s protocol. Luminescence was read and quantified using the FLUOSTAR Omega (BMG Labtech) plate reader.

### Statistical Analysis

All details of statistical analyses are outlined within the relevant figure legend, including test, replicates and any correction for multiple comparisons.

## Supporting information

Chammas_supp

## Data availability

RNA-seq data have been uploaded to GEO, accession GSE279214.

## Acknowledgements

We thank Lila Allou, Veronique Azuara, Vicki Metzis and members of the Percharde lab for project input and critical reading of the paper. We thank Yi Xuan Low for technical support with lentivirus generation. Thank you to Sanna Vuoristo for sharing iDUX4 hESCs. We acknowledge use of the Jex HPC cluster (Young and Cocktas 2023) at the MRC Laboratory of Medical Sciences and the resources and support provided by the IT and Bioinformatics Facilities. We are also grateful to LMS Microscopy, Flow cytometry and Genomics core facilities for all their assistance. Work in the Percharde lab is supported by a UKRI Future Leaders Fellowship (MC_EX_MR/X022560/1), a UKRI Impact Acceleration Award, and MRC funding (MRC; MC_UP_1605/4).

## Author Contributions

Conceptualization: MP, PC

Methodology: MP, PC, GY

Investigation: PC, LS, BJL, SQX,RTW, MHD

Visualization: PC, GY, DD, BJL, MP

Supervision: MP, MTM, MMK, GY

Writing—original draft: MP, PC

Writing—review & editing: MP, PC

