## Supplementary material for "CRISPRa-mediated disentanglement of the Dux-MERVL axis in the 2C-like state, totipotency and cell death": Chammas_supp

**List of data:**

Figure S1
Figure S2
Figure S3
Figure S4
Figure S5
Figure S6

Table S1: list of primers, sgRNAs and antibodies
Table S2: normalized gene and TE expression data

**A**

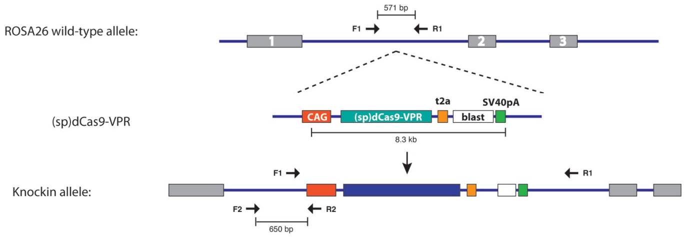


**B**

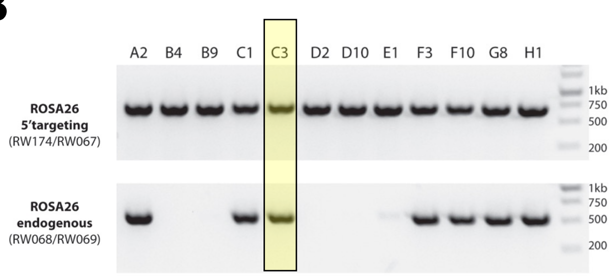


**C**

*MT2_Mm consensus*

```
>DF0004155:MT2_Mm
TGTAGTGGCTATTCCTGGTTGTCAACTTGACAATATTTGGAATGAACTACAATCCGGAATTGGAAGGCTCACCAGTG
ACCCTTATCTGGAGGCTTGGAGATCCTTATCTGGATCTTGGTTTGAAGATCTTGAGCCATAGTGGCTATGGATTCCA
GAAGATTGAATCTCCGAGTTTAAGGAACACACCTTTAATCTGGGCTACGCCTTTCATCTGGGATTAAAGGTGTGGTG
GAACACACCTTTAATCTGGGCTACACCTTCTGCTGGAGACAATATAAGGACATTGGAAGAAGGGAGTCTAGCTCTTG
CTCTTGCTCCTTCGCCTGCTTGCTGCGTGAGACTGAGTAACTGCTAGATCCTTGGACTTCCATTCACAGCTGCGACT
GAACAATTGTTGGGAATTGGGCTGCCGACTGTAAGTCATCAATAAATTCCTTTACTATCTAGAGACTATCCATAAGT
TCTGTGACTCTAGAGAACCCTGACTAATACA
```

**MERVL sgRNA 1**
**MERVL sgRNA 2**
**MERVL sgRNA 3**
**Mt2_Mm TSS**

**D**

*mDux*

```
CCCCGTGGCTTGCCAGGGATGAGGAGCCTTTGGGGAGATTTTGCTGAGTGTCAGAGGACGCTTGAGGTCCGCCTCCTGG
TGCCCTGATGTCAGGTGCCAGGCGTGGCAGGCTTGGCGCTTCTTGGGGGCAATGCGAGAAAGGGGTAGGCATGTTCTGT
AGCCCAAGGCGCCTAACTGGTAGGTAGTGGGCGGGACTACCTGAGCAGAGGCAGAGGTATTTAAGGGGCAGTGGTCACA
GCCACTCTGCTGGCAGTTGTTGCAGCTTGTGTTTGTTCTGAAGCTGTCCTGAGTCGAGTCTCCCAAGGTGAGGACTCCT
GGGAGGCCGTCATTGGCACCATG
```

**Dux sgRNA 1**
**Dux sgRNA 2**
**Dux coding start**

**Figure S1 dCAS9-VPR cell line and CRISPRa guide selection**

A) Diagram of *Rosa26* locus targeting strategy with *CAG-dCas9-t2a-Bsr* construct. Targeting was enhanced by co-transfection with a plasmid containing wt Cas9 plus sgRosa26 guide.
B) Genotyping results for dCas9-VPR ESCs. Clone 3, used for 2C-GFP/CD4 targeting, is highlighted.
C) Consensus sequence of the MERVL LTR, MT2_Mm, showing the positions of *sgMERVL1/2/3.*
D) Sequence of the mouse *Dux* promoter region, with the position of *sgDux1* and *sgDux2* indicated.

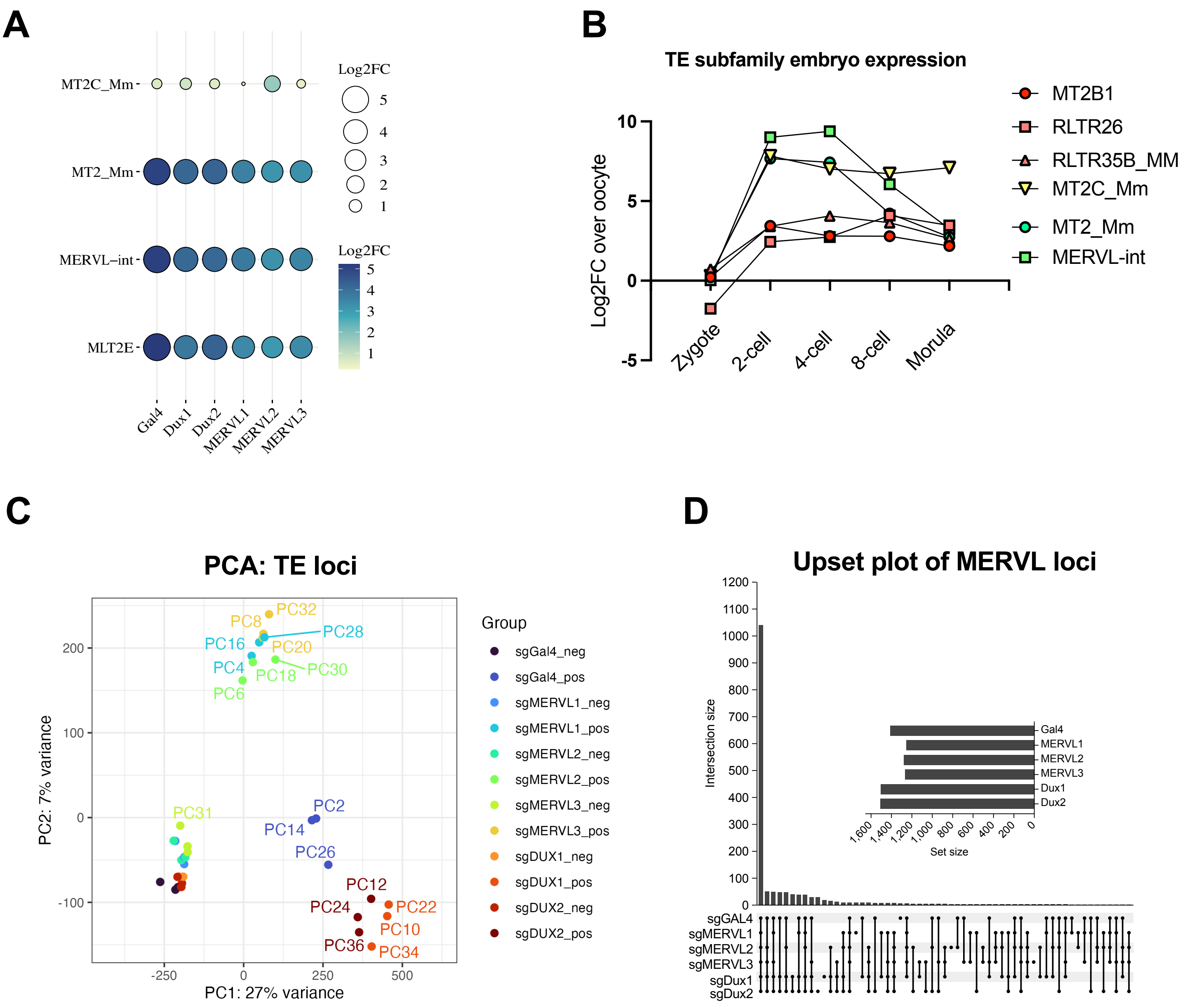


**Figure S2 RNA-seq analysis of TE subfamilies and TE loci**

A) Bubble plot showing the log2-fold change(FC) in expression in the indicated TE subfamilies in GFP-positive over GFP-negative samples.

B) Expression data showing log2FC of the indicated TE subfamilies in each embryo stage, relative to levels in oocytes. Data from *Modzelewski et al., 2021*, *Xue et al., 2013*

C) PCA plot of GFP-positive and negative samples clustered according to expression of individual TE loci. The first two principal components are shown.

D) Upset plot showing overlap in the number of MERVL (MT2_Mm, MERVL-int) loci significantly upregulated by each sgRNA in GFP-positive over negative samples (FDR < 0.05).

### Figure S3 Chammas et al.

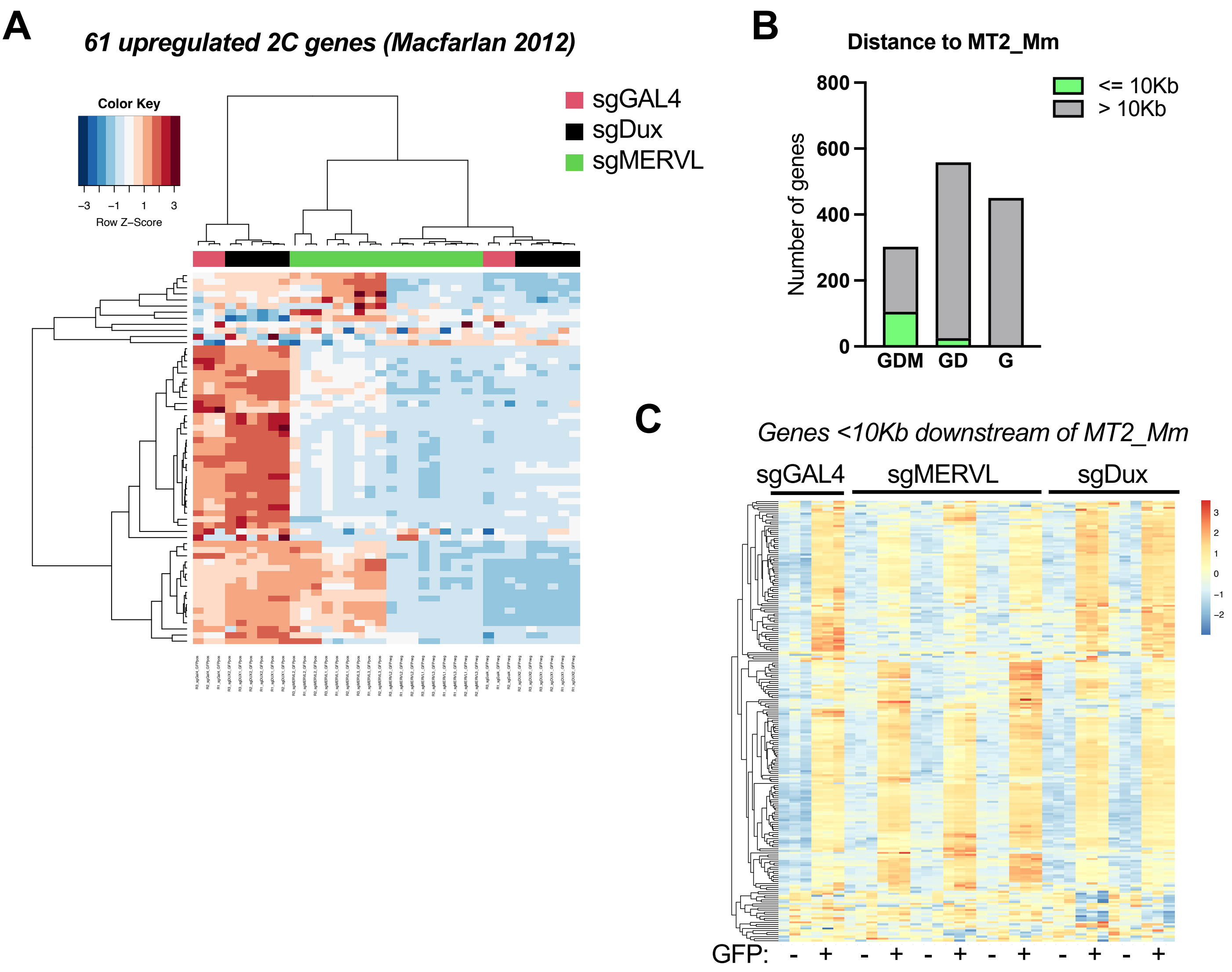


**Figure S3 RNA-seq analysis of gene expression data**

A) Heatmap showing the expression of 2C-specific genes (Macfarlan et al., 2012) across all GFP-positive and negative samples. Genes and samples are grouped by unsupervised hierarchical clustering.

B) Histogram showing the number of significantly upregulated genes in GDM, GD or G datasets that are within, or greater than, 10Kb from a MT2_Mm element.

C) Heatmap showing the expression of all genes within 10Kb downstream of an MT2_Mm element in GFP-positive and negative samples. Genes, but not samples, are grouped by unsupervised hierarchical clustering.

**Figure S4** **Chammas et al.**

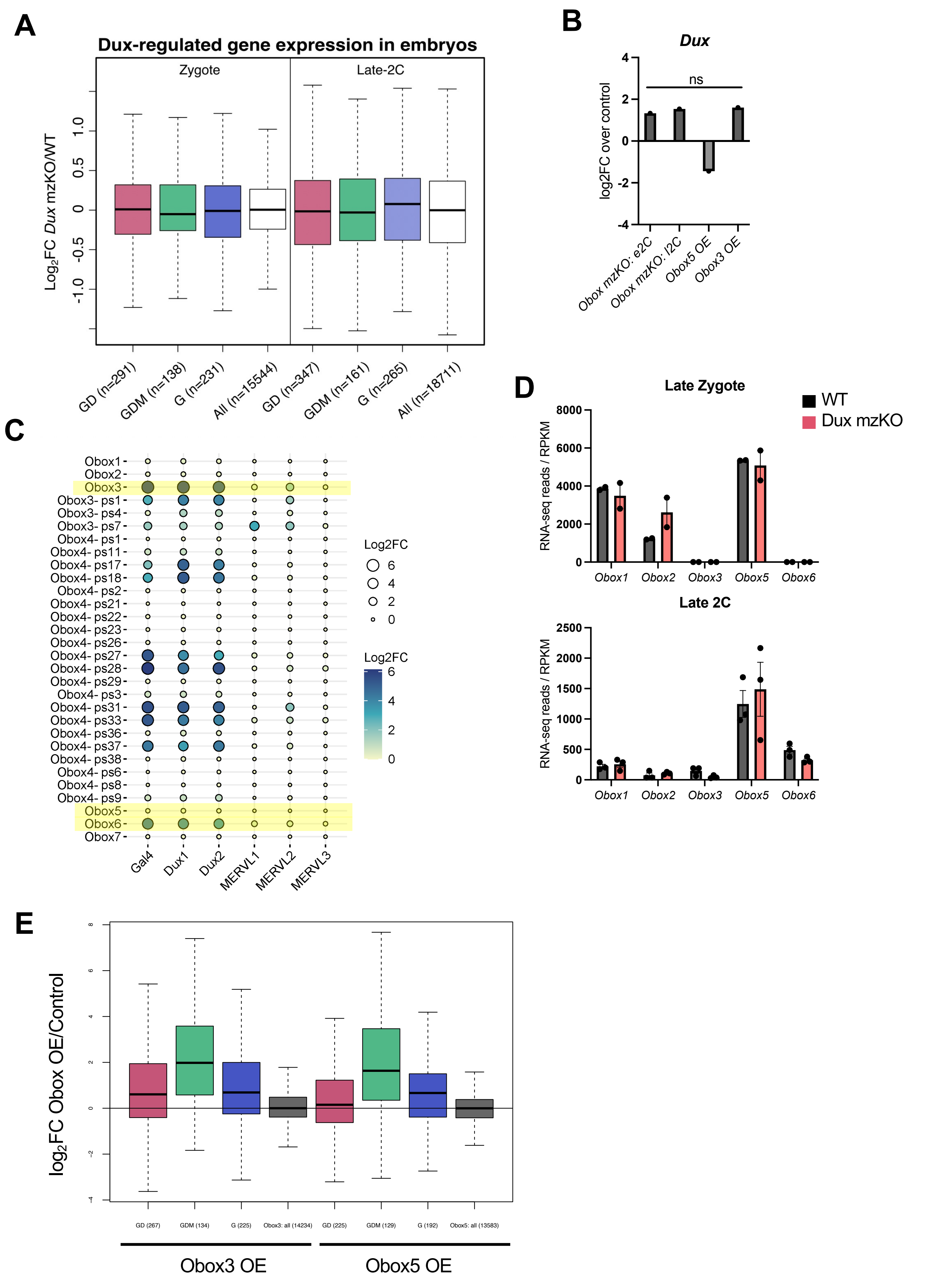

**Figure S4 RNA-seq analysis of gene expression data**

A) Boxplots showing log2FC of the indicated gene sets (GD, GDM, G, all) in Dux mzKO over WT embryos. Gene sizes are shown in brackets. Data from Chen et al., 2019

*B)* *Dux* expression in Obox depleted or overexpressing samples (embryos or ESCs, respectively). Data from Ji et al., 2023, ns = not significant (FDR >0.05)

C) Bubble plot showing log2FC in the indicated *Obox* transcripts in GFP-positive over negative samples from each sgRNA transfection.

D) Histogram showing the normalized expression (RPKM) of *Obox* factors in WT versus *Dux* mzKO embryos. Data from Chen et al., 2019

E) Boxplots showing log2FC of the indicated gene sets (GD, GDM, G, all) in ESCs upon either Obox3 or Obox5 overexpression. Gene sizes are shown in brackets. Data from Ji et al., 2023

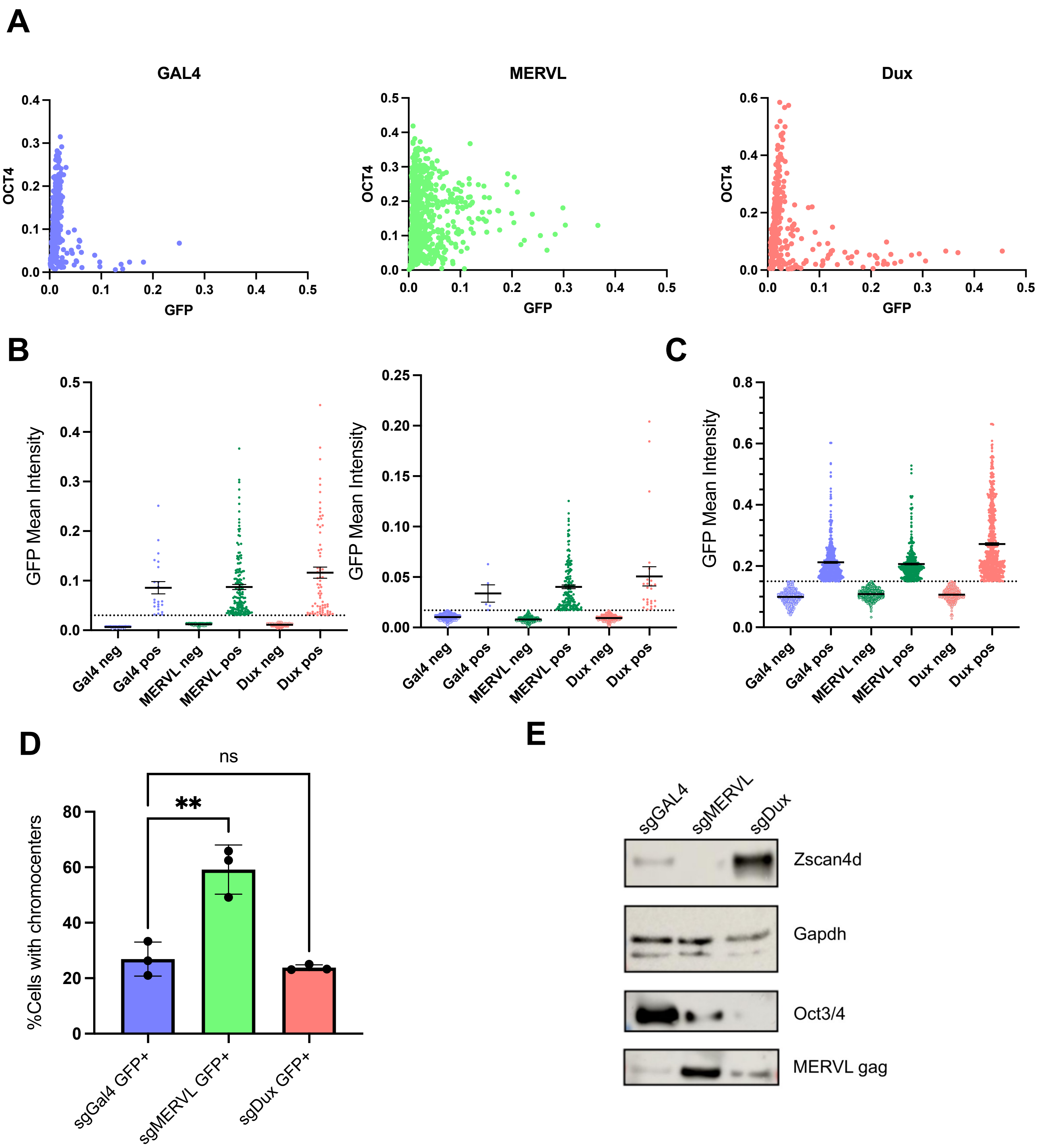


**Figure S5**

A) Mean intensity of OCT4 and GFP in MACS-sorted cells after indicated sgRNA transfections. Data are combined from 2 independent experiments.

B-C) GFP mean intensity in cells from MACS purifications, post thresholding (dotted line) to remove outliers based on GFP. Data in B) relate to Oct4 IFs – combined from 2 experiments and in (C) to nucleolar circularity – 1 representative experiment.

D) Quantification of the percentage of GFP-positive cells from the indicated sgRNA transfections that are chromocenter positive. Data are from 3 independent experiments. $P < 0.01$, one-way ANOVA with Dunnett's multiple comparisons test.

E) Western blot of samples from unsorted CVG ESCs 72h after transduction with the indicated sgRNAs. Representative of n=2-3 experiments.

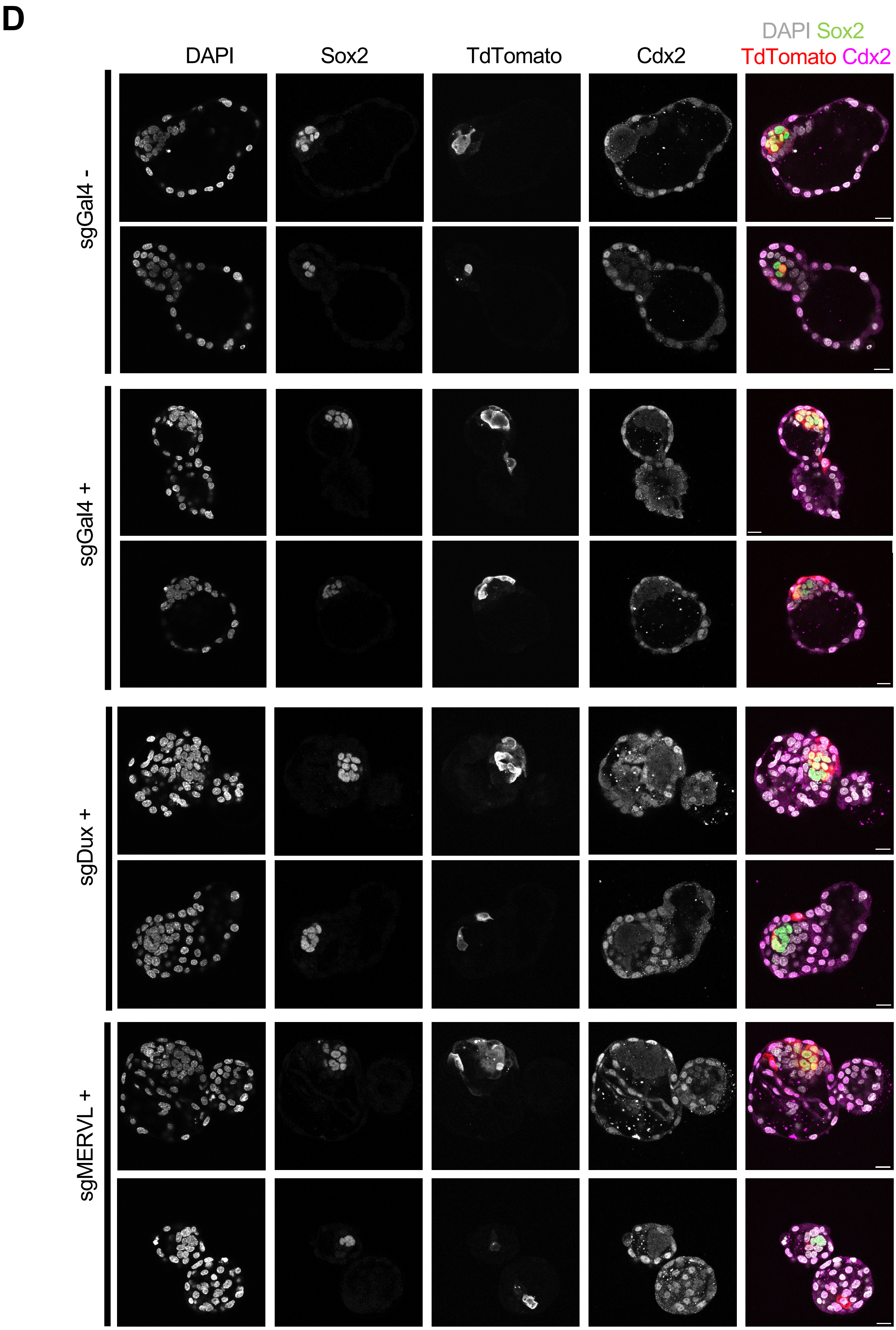


**Figure S5 (cont'd)**

F) Representative confocal images of embryonic day (E) 4.5 blastocysts following injection of sgGal4 GFP-, sgGal4 GFP+, MERVL GFP+ or Dux GFP+ cells at the 8-cell stage. Embryos were immunostained for Sox2, Cdx2, and TdTomato and counterstained with DAPI. Two examples are shown for each condition. For each of sgDux and sgMERVL conditions, one example is the same embryo as shown in Figure 4F. Maximum projection of 5 z-sections of 1 micron. Scale bar = 20um.

### Figure S6

Chammas et al.

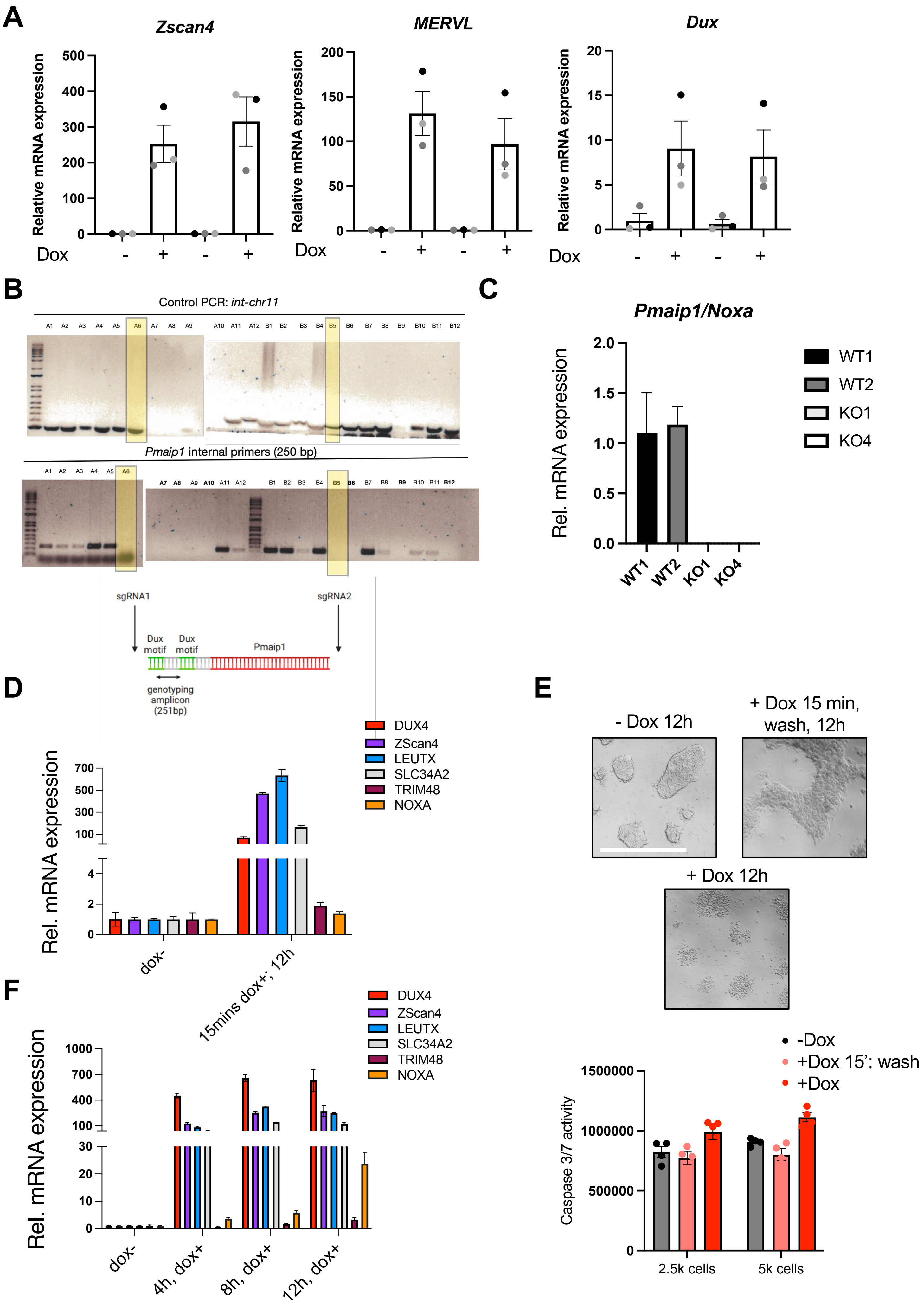

**Figure S6 NOXA analysis in mouse and human cells**

A) RT-qPCR analysis of the indicated 2C genes in iDux ESCs 24h after dox treatment. Data are mean +/- s.e.m, n=3 independent experiments.
B) Genotyping PCR strategy and results for the generation of *Noxa* KO mouse ESCs
C) RT-qPCR validation showing no expression of *Noxa* in KO ESCs. Data are mean +/- s.e.m of n=3 independent experiments.
D) qRT-PCR in human H9 iDUX4 hESCs for the indicated genes. Samples were harvested 12 hours after a 15 minute pulse induction with Doxycycline (dox) to induce DUX4 expression. Data are mean +/- s.e.m of n=3 wells.
E) Representative images and caspase-3/7 assay of iDUX4 hESCs in the absence or presence of Dox for a 15 min pulse or kept on cells constantly. Data are representative of 3 experiments, harvested at 12-14h after Dox. Scale bar, 100 um
F) qRT-PCR expression in H9 iDUX4 hESCs for the indicated genes following increasing duration of continuous Doxycycline-mediated DUX4 induction. Data are mean +/- s.e.m of n=3 wells.
